# Simultaneous, cell-intrinsic downregulation of PD-1 and TIGIT enhances the effector function of CD19-targeting CAR T cells and promotes an early-memory phenotype

**DOI:** 10.1101/2020.11.07.372334

**Authors:** Young-Ho Lee, Hyeong Ji Lee, Hyung Cheol Kim, Yujean Lee, Su Kyung Nam, Cedric Hupperetz, Jennifer SY Ma, Xinxin Wang, Oded Singer, Won Seog Kim, Seok Jin Kim, Youngil Koh, Inkyung Jung, Chan Hyuk Kim

**Affiliations:** Department of Biological Sciences, Korea Advanced Institute of Science and Technology, Daejeon 34141, Republic of Korea; Curocell Inc., Daejeon 34109, Republic of Korea; California Institute for Biomedical Research, 11119 North Torrey Pines Road, La Jolla, CA 92037, USA; Division of Hematology and Oncology, Department of Medicine, Samsung Medical Center, Sungkyunkwan University School of Medicine. 06351, Republic of Korea; Department of Internal Medicine, Seoul National University Hospital, Seoul 03080, Republic of Korea

**Author notes:** These authors contributed equally to this work (Young-Ho Lee, Hyeong Ji Lee). Correspondence to Chan Hyuk Kim.

## Abstract

CD19-targeting chimeric antigen receptor (CAR) T cells have become an important therapeutic option for patients with relapsed and refractory B cell malignancies. However, recent clinical data indicate that a significant portion of patients still do not benefit from the therapy owing to various resistance mechanisms, including high expression of multiple inhibitory immune checkpoint receptors on activated CAR T cells. Here, we report a lentiviral two-in-one CAR T approach in which two checkpoint receptors are downregulated simultaneously by a dual short-hairpin RNA (shRNA) cassette integrated into a CAR vector. Using this system, we evaluated CD19-targeting CAR T cells in the context of four different checkpoint combinations—PD-1/TIM-3, PD-1/LAG-3, PD-1/CTLA-4 and PD-1/TIGIT—and found that CAR T cells with PD-1/TIGIT downregulation uniquely exerted synergistic antitumor effects in mouse xenograft models compared with PD-1 single downregulation, and maintained cytolytic and proliferative capacity upon repeated antigen exposure. Importantly, functional and phenotypic analyses of CAR T cells as well as analyses of transcriptomic profiles suggested that downregulation of PD-1 enhances short-term effector function, whereas downregulation of TIGIT is primarily responsible for maintaining a less-differentiated/exhausted state, providing a potential mechanism for the observed synergy. The PD-1/TIGIT–downregulated CAR T cells generated from diffuse large B-cell lymphoma patient-derived T cells using a clinically applicable manufacturing process also showed robust antitumor activity and significantly improved persistence *in vivo* compared with conventional CD19-targeting CAR T cells. Overall, our results demonstrate that the cell-intrinsic PD-1/TIGIT dual downregulation strategy may prove effective in overcoming immune checkpoint-mediated resistance in CAR T therapy.

## Introduction

CD19-specific chimeric antigen receptor (CAR) T cell therapy has proven highly effective in the treatment of relapsed or refractory (R/R) B cell malignancies such as acute lymphoblastic leukemia (ALL) and non-Hodgkin lymphomas (NHLs)^1–6^. However, recent studies have reported that about half of treated patients do not benefit from the treatment^2–5^, and that chronic lymphoblastic leukemia (CLL) patients show only a marginal response^7^. Treatment failure is often attributed to the loss of CD19 expression in cancer cells^8^, but can also involve the limited expansion, poor persistence, and dysfunctional state of infused CAR T cells^9–11^.

T cells express inhibitory immune checkpoint receptors (ICRs) that are essential for maintaining peripheral tolerance and regulating immune responses^12,13^. However, in chronic inflammatory conditions and cancer, persistent ICR expression leads to dysfunction of antigen-specific T cells which includes reduced cytolytic activity, cytokine secretion, survival, and proliferation^12,14–16^. Consistent with this, monoclonal antibody-mediated blockade of either cytotoxic T-lymphocyte-associated protein 4 (CTLA-4) or programmed cell death protein 1 (PD-1) is effective against many cancers^15,17^. However, single ICR blockade therapy only yields a sustained response in a small subset of patients and often fades over time^18^. Instead, recent murine and human studies have shown that simultaneous or sequential blockade of multiple ICRs can have synergistic effects on disease control^15,19–24^.

In the context of CAR T therapy, previous preclinical and clinical studies have evaluated the co-administration of PD-1-blocking antibody, demonstrating enhanced antitumor activity and *in vivo* expansion of CAR T cells^25–27^. Genetically, CAR T cells have been modified to secrete single-chain variable fragments (scFv) against PD-1^28^ and to express dominant-negative^26,29^ or chimeric-switch receptors^30–32^. In addition, CRISPR/Cas9-based knockout and short-hairpin RNA (shRNA)-based knockdown have been used to achieve sustained downregulation of ICRs in a cell-intrinsic manner^26,33–38^. However, most of these genetic approaches target a single immune checkpoint receptor, and few studies have examined the therapeutic effects and cellular mechanisms of CAR T cells with multiple cell-intrinsic ICR blockade^39^.

Here, we describe a dual shRNA-based approach that allows for the simultaneous downregulation of two ICRs in CAR T cells. With this, we found that different combinations of ICR downregulation differentially affected the activity of CAR T cells, with some being detrimental. In particular, the downregulation of PD-1 and TIGIT (T-cell immunoreceptor with Ig and ITIM domains) was the most effective, significantly enhancing the efficacy of CAR T cells generated from a healthy donor and NHL patient T cells. Importantly, RNA sequencing and cellular analysis suggested that the downregulation of PD-1 and TIGIT in CAR T cells distinctly affect their effector function and differentiation, which likely accounts for the synergistic improvement in activity compared to the downregulation of PD-1 or TIGIT alone.

## Results

### PD-1 downregulation using a two-in-one lentiviral vector system enhances the antitumor activity of CD19-targeting CAR T cells

Given that PD-1 is one of the major ICRs, we first designed a two-in-one lentiviral vector that expresses shRNA against PD-1 alongside a 41BB-based 2^nd^-generation CD19-targeting CAR (**Fig. 1a**). We tested 3 different PD-1-targeting *shRNAs (shRNA* #1-3 **Supplementary Fig. 1a**) and 3 different Pol III promoters (mouse U6, human U6, and human H1)^40^ and found that the 19PBBz construct expressing *shPD-1* #1 driven by mouse U6 efficiently downregulated PD-1 (**Fig. 1b and d)** and had no negative effect on CAR expression (**Supplementary Fig. 1b**) or CAR T cell expansion (**Fig. 1c and e**). We next cocultured 19PBBz CAR T cells with CD19-positive target cells with or without PD-L1 expression (Nalm-6, Nalm-6-PD-L1, K562-CD19, and K562-CD19-PD-L1 cells; **Supplementary Fig. 2**) and found that 19PBBz CAR T cells exhibited enhanced cytotoxic activity (**Fig. 1f**) and proliferation (**Fig. 1g**) compared to control CAR T cells (19GBBz and 19BBz, with and without the mU6-*shGFP* cassette, respectively), which depended on target PD-L1 expression. In addition, 19PBBz cells exhibited improved *in vivo* antitumor activity in mouse xenograft models of disseminated human blood cancer (Nalm-6-GL-PD-L1 and Nalm-6-GL) in a manner dependent on target PD-L1 expression (**Fig. 1h and Supplementary Fig. 3**). Importantly, PD-1 expression levels in CAR T cells isolated from mice on day 43 indicated that our two-in-one vector system persistently downregulated PD-1 under *in vivo* conditions (**Fig. 1i**).

**Figure 1.**
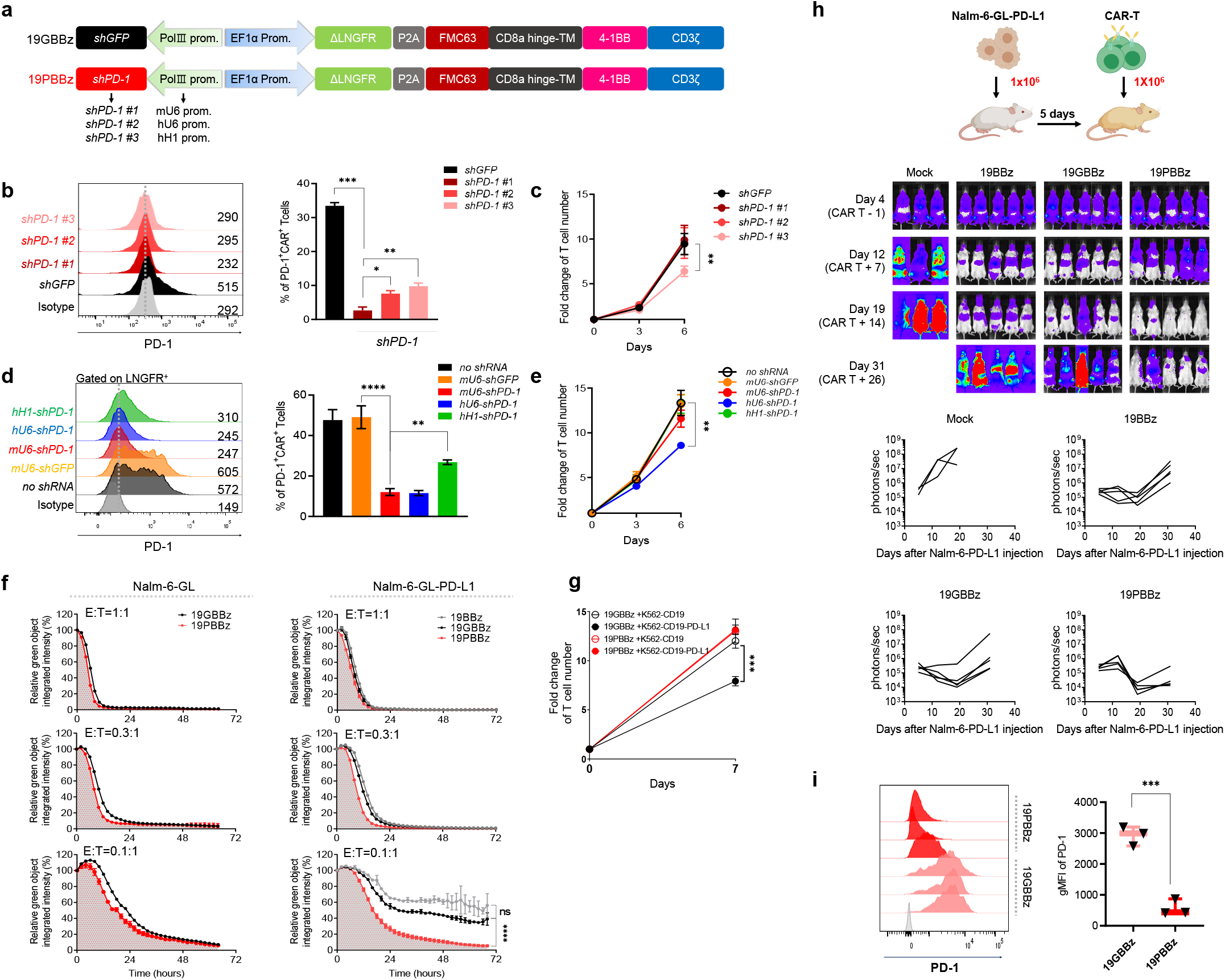
PD-1 downregulation enhances the antitumor function of CAR T cells. **(a)** Schematic representation of the lentiviral two-in-one vector carrying a CD19-CAR and shRNA expressing module. **(b)** PD-1 expression levels of CAR T cells with different shPD-1 candidates as determined by flow cytometry on day two after stimulation with γ-irradiated Nalm-6-GL cells. Gray denotes the isotype control. Data are the pooled mean ± SD from three independent experiments, each performed in triplicates. **(c)** Cell counts from the homeostatic expansion of LNGFR^+^ CAR T cells with PD-1 downregulation candidates on days 3 and 6 after cell seeding. Data are the pooled mean ± SD from three independent experiments performed in triplicates. **(d)** The effects of hH1-, hU6-, and *mU6-shPD1* on PD-1 expression was analyzed by flow cytometry two days after stimulation with γ-irradiated K562-CD19 cells. Data are the pooled mean ± SD from two independent experiments performed in triplicates. **(e)** Cell counts from the homeostatic expansion of CAR T cells with each Pol III promoter after LNGFR^+^ isolation on days 3 and 6 after cell seeding. Data are the pooled mean ± SD from two independent experiments performed in triplicates. **(f)** CAR T cells were incubated with GFP-expressing Nalm-6-GL or Nalm-6-GL-PD-L1 cells at a 1:1, 0.3:1, and 0.1:1 effector: target (E:T) ratio. GFP intensity was measured every two hours using the IncuCyte S3 live-cell imaging system. The relative percentage of total integrated GFP intensity was calculated as [GFP intensity at each time point / GFP intensity at 0 h]*100. Representative mean ± SD from two independent experiments performed in triplicates. **(g)** CAR T cells were incubated with γ-irradiated K562-CD19 or K562-CD19-PD-L1 cells at a 1:3 E:T ratio and counted on day 7. Data are the pooled mean ± SD from two independent experiments performed in triplicates. **(h)** NSG mice were injected intravenously with 1×10^6^ Nalm-6-GL-PD-L1 leukemia cells. 5 days later, 1×10^6^ CAR T cells were injected intravenously. Tumor burden was monitored based on the bioluminescence intensity from the IVIS imaging system. Data are from n = 3 mock and n = 5 19BBz, 19GBBz, and 19PBBz mice, respectively, **(i)** PD-1 expression levels of CAR T cells from Nalm-6-GL-PD-L1-bearing mice at day 43. Gray denotes isotype control. Data are the mean ± SD from three mice per group. Statistical analysis was done by One-Way ANOVA for (b-f) and unpaired two-tailed t-test for (g and i). *p < 0.05, ** p < 0.01, *** = p < 0.001, **** = p < 0.0001, ns = not significant.

After optimizing the shRNA expression cassette targeting PD-1, we wondered if this system has differential effects depending on the costimulatory domain within the CAR. Because CD28 has been shown as the major target of PD-1^41^, we also generated a 19P28z construct that replaces the 41BB costimulatory domain in 19PBBz with CD28 and confirmed similar CAR expression levels between 19P28z, 19PBBz, and their controls (19G28z and 19GBBz, **Supplementary Fig. 4a**). We then confirmed that, as 19PBBz, 19P28z exhibited reduced PD-1 protein expression compared to its control upon stimulation with γ-irradiated K562-CD19 (**Fig. 2a**). When we evaluated the *in vitro* cytotoxicity, we found similarly improved cytotoxicity of freshly-prepared 19P28z and 19PBBz CAR T cells compared to their controls against Nalm-6-GL-PD-L1 cells (**Fig. 2b**). However, when we compared the cytotoxicity of CAR T cells that were stimulated with Nalm-6-PD-L1-CD80 for 6 days, we found that 19P28z CAR T cells showed reduced cytotoxicity compared to 19PBBz CAR T cells (**Fig. 2c**).

**Figure 2.**
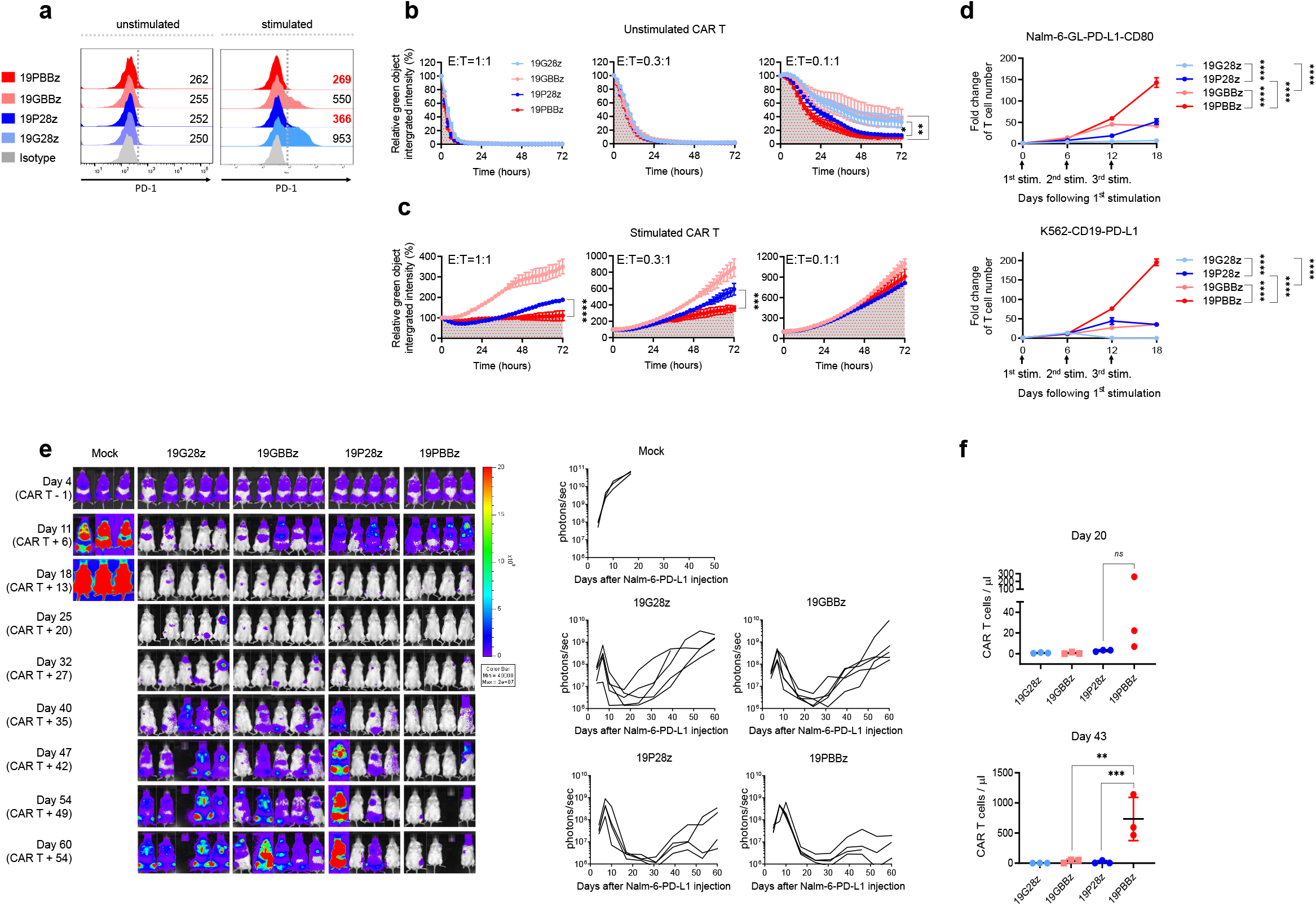
CAR costimulatory domains differentially affect the *in vivo* persistence of PD-1-downregulated CAR T cells. **(a)** PD-1 expression level of CAR T cells with CD28 or 41BB costimulatory domains two days after stimulation with γ-irradiated K562-CD19 cells. Numbers denote the gMFI of PD-1. **(b)** Unstimulated and **(c)** stimulated CAR T cells were incubated with Nalm-6-GL-PD-L1 cells at a 1:1,0.3:1,0.1:1 E:T ratio and analyzed using the IncuCyte S3 system. Stimulated CAR T cells were generated by coincubation with Nalm-6-PD-L1-CD80 cells for 6 days prior to cytotoxicity assay. Data are the representative mean ± SD from two independent experiments performed in triplicates. **(d)** CAR T cells were incubated with γ-irradiated Nalm-6-GL-PD-L1-CD80 or K562-CD19-PD-L1 cells at 1:3 effector: target (E: T) ratio and counted on day 6 after each stimulation. Data are the mean ± SD from two independent experiments performed in triplicates. **(e)** NSG mice were injected intravenously with 1×10^6^ Nalm-6-GL-PD-L1 leukemia cells. 5 days later, 1×10^6^ CAR T cells were injected intravenously. Tumor burden was monitored based on the bioluminescence intensity from the IVIS imaging system. Data are from n = 3 mock, n = 5 19G28z and 19GBBz, and n = 4 19P28z and 19PBBz. **(f)** The number of CAR T cells in mouse blood was determined on day 20 and 43 after CAR T cell injection. Data are mean ± SD from three mice per group. Statistical analysis for (a-d and f) was done by One-Way ANOVA. * = p < 0.05, ** = p < 0.01, *** = p < 0.001, **** = p < 0.0001, ns = not significant.

Additionally, 19PBBz was the only construct capable of robust proliferation upon repeated stimulation (**Fig. 2d**). Consistent with these *in vitro* observations, 19PBBz CAR T cells demonstrated superior *in vivo* antitumor activity compared to 19P28z CAR T cells in mouse xenograft models of disseminated human blood cancer (Nalm-6-GL-PD-L1) and exhibited sustained, high-level accumulation of CAR T cells in the blood for up to 43 days (**Fig. 2e and f**).

Collectively, our results show that the modification of the lentiviral vector to express an additional shRNA cassette that specifically targets PD-1 is a viable approach for enhancing the antitumor activity of CD19-targeting CAR T cells against PD-L1-positive cancer cells. In addition, we observed that this system had superior activity with a 41BB costimulatory domain compared to CD28, exhibiting a delayed exhaustion phenotype and longer persistence *in vivo.*

### Simultaneous downregulation of PD-1 and TIGIT further enhances the *in vivo* functionality of CD19-targeting CAR T cells

Although PD-1/PD-L1 blockade has been successful as a monotherapy in R/R Hodgkin lymphoma (HL), responses have been modest in R/R diffuse large B-cell lymphoma (DLBCL) and relapsed chronic lymphocytic leukemia (CLL) patients^42–45^. Furthermore, multiple ICRs are frequently expressed on intratumoral T cells from NHL patients^46^, and PD-1/PD-L1 blockade is known to induce compensatory upregulation of alternative ICRs^19,47^.

Therefore, we reasoned that simultaneous downregulation of additional ICRs would further enhance the antitumor activity of PD-1-downregulated CAR T cells. We chose four well-known ICRs: T cell immunoglobulin and mucin domain-containing protein 3 (TIM-3), lymphocyte-activation gene 3 (LAG-3), TIGIT, and CTLA-4, and confirmed that they are expressed on activated T cells *in vitro,* albeit with different kinetics and to different extents (**Supplementary Fig. 5**). After selecting shRNA sequences that individually downregulated their indicated receptors without affecting CAR expression (**Supplementary Figs. 6–8**), we modified our two-in-one lentiviral vector to include a second shRNA cassette (**Fig. 3a**) and generated CD19-targeting dual-shRNA CAR vectors with four ICR combinations: PD-1/TIM-3, PD-1/LAG-3, PD-1/TIGIT and PD-1/CTLA-4. We confirmed that PD-1 downregulation was similar across all constructs, and that the secondary shRNAs also specifically downregulated their respective ICRs without affecting CAR expression **(Fig. 3b, Supplementary Fig. 9a and b)**. To assess the effect of dual ICR downregulation on antitumor activity, we injected 1×10^6^ CAR T cells into mice bearing Nalm-6-GL-PD-L1 leukemia cells that express ligands for TIGIT (CD112/155), LAG-3 (HLA-DR), and TIM-3 (galectin-9), but not CTLA-4 (CD80) (**Supplementary Fig. 2 and 9c-e**). Unexpectedly, we found that three of the four combinations (PD-1/TIM-3, PD-1/LAG-3, and PD-1/CTLA-4) failed to improve the *in vivo* efficacy of CD19-targeting CAR T cells compared with single downregulation of PD-1; in the case of PD-1/LAG-3, dual downregulation severely compromised antitumor activity **(Fig. 3c,d and Supplementary Fig. 10a)**. However, the PD-1/TIGIT combination reduced the frequency of leukemia relapse and improved the survival of mice compared to single PD-1 downregulation **(Fig. 3c and d)**. Further, dual PD-1/TIGIT downregulation outperformed single PD-1 downregulation in a stress test with two lower doses (0.5×10^6^ and 0.25×10^6^) of CAR T cells in the same mouse model (**Fig. 3e and f and Supplementary Fig. 10b**).

**Figure 3.**
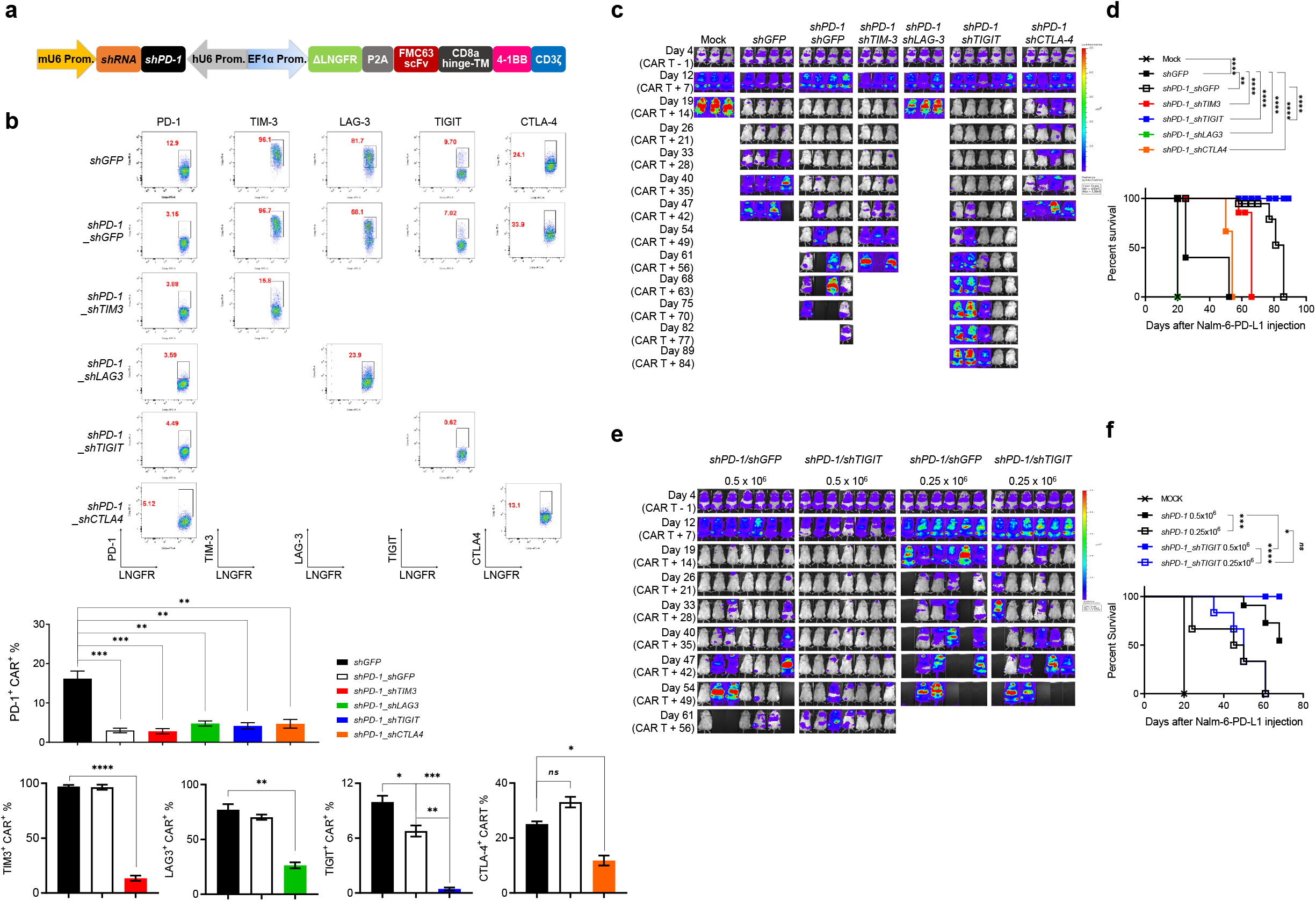
Simultaneous downregulation of PD-1 and TIGIT further enhances the *in vivo* functionality of CD19-targeting CAR T cells. **(a)** Schematic representation of the engineered two-in-one vector system carrying dual shRNA cassettes for two ICRs. **(b)** Dual downregulation efficiency of each ICR in CAR T cells stimulated for 48 hours with γ-irradiated K562-CD19 cells. FACS plots are representative data from two independent experiments performed in duplicates and the bar graphs are the pooled mean ± SD. **(c)** NSG mice were injected intravenously with 1×10^6^ Nalm-6-GL-PD-L1 leukemia cells. 5 days later, 1×10^6^ CAR T cells with each dual downregulation *(shGFP, shPD-1/shGFP, shPD-1/shTIM-3, shPD-1/shLAG-3, shPD-1/shTIGIT, shPD-1/shCTLA-4)* were injected intravenously. Tumor burden was monitored based on the bioluminescence intensity from the IVIS imaging system. Data are from n = 3 mock, *shPD-1/shTIM-3* and *shPD-1/shLAG-3,* n = 4 *shGFP, shPD-1/shGFP,* and *shPD-1/shCTLA-4,* n = 5 *shPD-1/shTIGIT.* **(d)** Kaplan-Meier survival analysis with the Log-rank (Mantel-Cox) test comparing each CAR T treated mice from (c). **(e)** NSG mice were injected intravenously with 1×10^6^ Nalm-6-GL-PD-L1 leukemia cells. 5 days later, 0.5×10^6^ or 0.25×10^6^ CAR T cells with PD-1 *(shPD-1/shGFP)* or PD-1/TIGIT *(shPD-1/shTIGIT)* downregulation were injected intravenously. Tumor burden was monitored based on the bioluminescence intensity from the IVIS imaging system. Data are from n = 7 mice for the 0.5×10^6^ dose groups and n = 6 mice for the 0.25×10^6^ dose groups. **(f)** Kaplan-Meier survival analysis with Log-rank (Mantel-Cox) test comparing CAR T treated mice from (e). Statistical analysis for (b) was done by One-Way ANOVA. * = p < 0.05, ** = p < 0.01, *** = p < 0.001, **** = p < 0.0001 ns, not significant.

**Figure 4.**
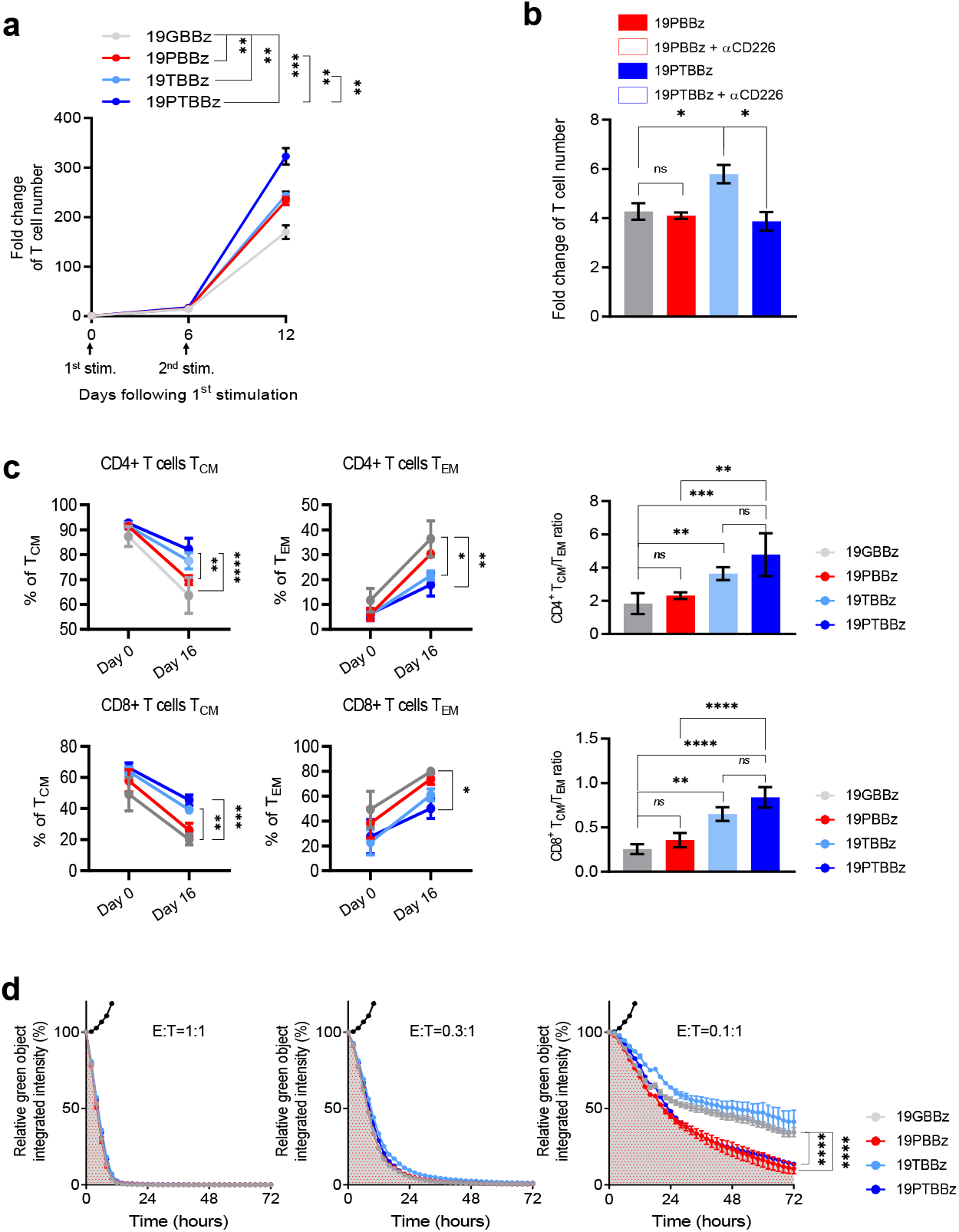
Downregulation of PD-1 and TIGIT distinctly affects the *in vitro* function of CAR T cells. **(a)** 1×10^6^ CAR T cells with single- or dual-downregulation were incubated with 3×10^6^ γ-irradiated Nalm-6-GL-PD-L1-CD80 cells every six days (1^st^ and 2^nd^ stimulation) and counted on day 6 after each respective stimulation. Data are the pooled mean ± SD from two independent experiments performed in triplicates. **(b)** 1×10^6^ 1^st^ stimulated day 6 19PBBz or 19PTBBz were incubated with 3×10^6^ γ-irradiated Nalm-6-GL-PD-L1-CD80 cells with or without 10 μg/mL CD226 blockade antibody for six days and counted. Data are the mean ± SD from one experiment performed in triplicates. **(c)** The expression of CD45RO and CCR7 was measured to distinguish the differentiation state of CD4^+^ and CD8^+^ 19GBBz, 19PBBz, 19TBBz, or 19PTBBz cells on day 16 (10 days after 2^nd^ stimulation). Data are the mean ± SD from 3 donors. T_CM_: CD45RO+CCR7+, T_EM_: CD45RO+CCR7-. **(d)** IncuCyte-based cytotoxicity kinetics of 1^st^ stimulation CAR T cells against Nalm-6-GL-PD-L1 cells on day 6 at the indicated ratios. Data are the representative mean ± SD from two independent experiments perfomed in triplicates. Filled-black dots represent co-culture with untranduced T cells. Statistical analysis for (a,b,c) was done by One-Way ANOVA and for (d) by unpaired two-tailed t-test. ** = p < 0.01, *** = p < 0.0001, **** = p < 0.0001. ns, not significant.

**Figure 5.**
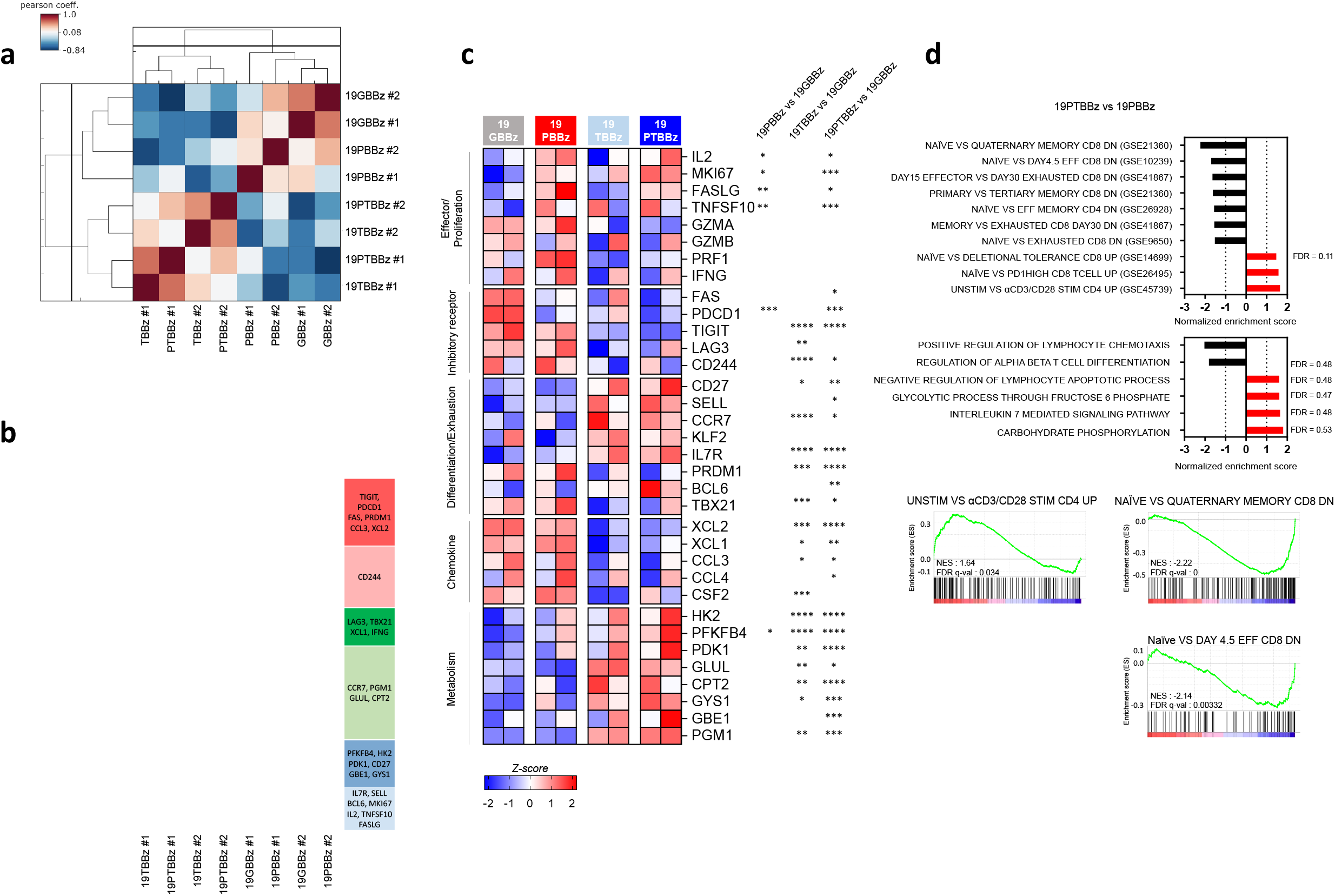
RNA-seq analysis uncovers the distinct roles of PD-1 and TIGIT downregulation. 2^nd^-stimulated 19GBBz, 19PBBz, 19TBBz, and 19PTBBz cells were prepared as shown in Supplementary Fig. 16 (a) for RNA-seq. **(a)** Pearson’s correlation analysis of the transcriptomic profiles and **(b)** hierarchical clustering of differentially-expressed genes from 2^nd^-stimulated 19GBBz, 19PBBz, 19TBBz, and 19PTBBz cells derived from two donors (FDR *q ≤* 0.1). **(c)** Heat map of selected genes associated with T cell function in 2^nd^-stimulated 19GBBz, 19PBBz, 19TBBz, and 19PTBBz cells. Asterisks represent the statistical significance as measured by *q*-value. **(d)** Normalized Enrichment Scores (NESs) of significantly enriched gene sets associated with phenotypic and functional T cell signatures in 2^nd^-stimulated 19GBBz, 19PBBz, 19TBBz, and 19PTBBz cells as determined by GSEA analysis. For all gene sets, FDR *q ≤* 0.03 unless otherwise indicated. * = *q* < 0.05, ** = *q* < 0.01, *** = *q* < 0.001, **** = *q* < 0.0001.

**Figure 6.**
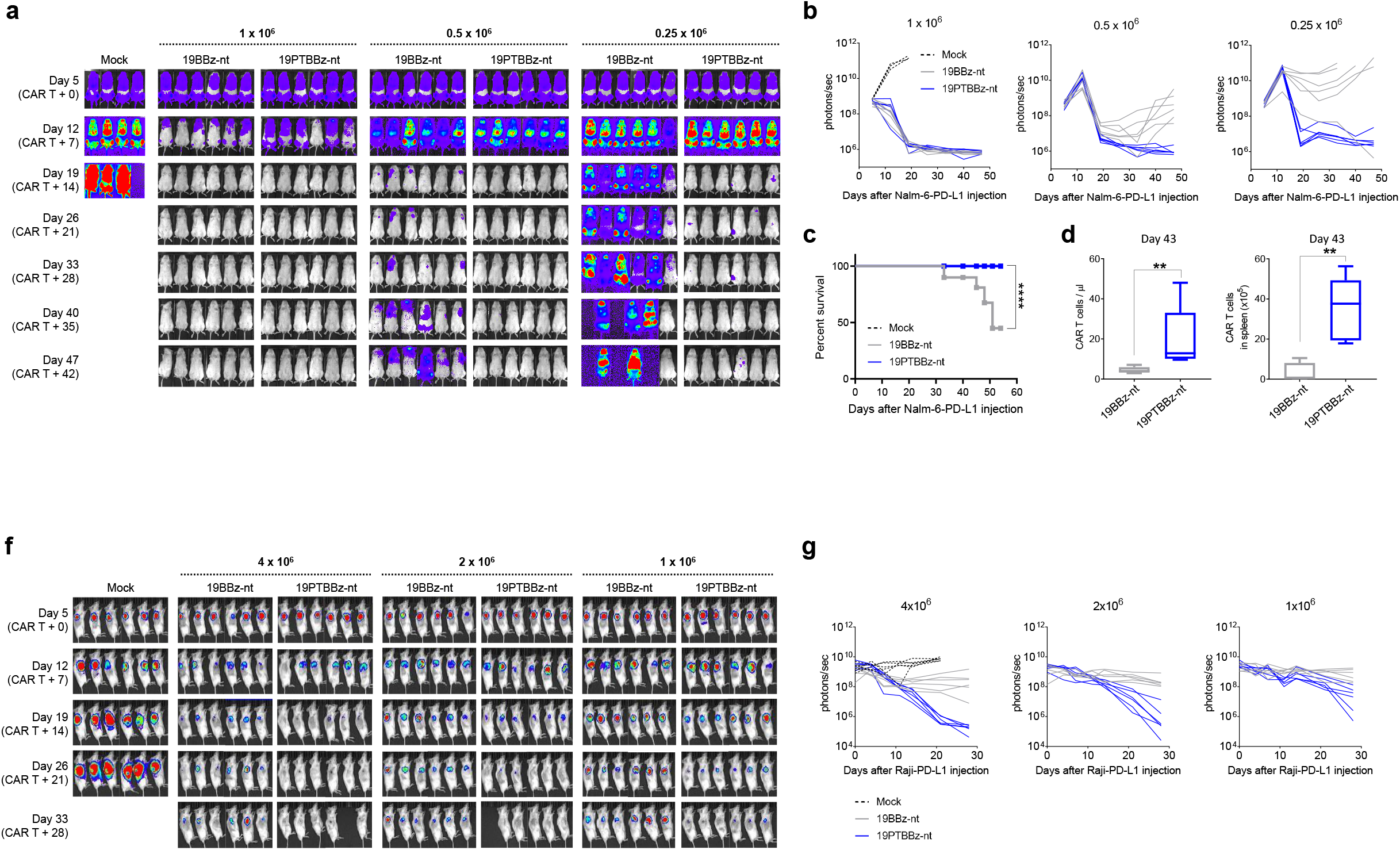
Clinical-scale manufactured CAR T cells with PD-1/TIGIT downregulation showed a superior in vivo functionality against leukemia and lymphoma tumor models. **(a,b)** NSG mice were injected intravenously with 1×10^6^ Nalm-6-GL-PD-L1 cells. 5 days post-tumor growth, Mock, 19BBz-nt, and 19PTBBz-nt cells were intravenously injected at the indicated doses. Tumor burden was monitored based on the bioluminescence intensity from the IVIS imaging system. Data are from n = 4 mock mice and n = 6 mice for all CAR T cell treatment groups. **(c)** Kaplan-Meier survival analysis with Log-rank test comparing the CAR T cell-treated mice at 0.25 x 10^6^ dose from (a,b). **(d)** The number of CAR T cells in the blood and spleen was determined 43 days after CAR T cell injection at a 0.5×10^6^ dose. Data are the mean ± SD from six mice per group. **(f-g)** NOG mice were injected subcutaneously on the right flank with 5×10^6^ Raji-GL-PD-L1 lymphoma cells. When the mean tumor volume reached approximately 100 mm^3^, CAR T cells were injected intravenously at the indicated doses. Tumor burden was monitored based on the bioluminescence intensity from the IVIS imaging system. Statistical analysis from (d) was done by unpaired two-tailed t-test. ** = p < 0.01, **** = p < 0.0001. nt = not tagged.

These results suggest that the effects of simultaneous cell-intrinsic downregulation of two ICRs can vary depending on the receptor combinations, with some being detrimental. Notably, dual downregulation of PD-1/TIGIT enhances the *in vivo* antitumor activity of CD19-targeting CAR T cells compared to single downregulation of PD-1.

### Downregulation of PD-1 and TIGIT distinctly affects the *in vitro* function of CAR T cells

To investigate the mechanistic basis of the cell-intrinsic PD-1/TIGIT dual downregulation effect, we generated CD19-targeting CAR T cells with single (19PBBz and 19TBBz) or dual (19PTBBz) downregulation of each receptor in parallel (**Supplementary Fig. 11**). An evaluation of the proliferative capacity of CAR T cells under repeated antigen exposure using γ-irradiated Nalm-6-PD-L1-CD80 cells showed that 19PTBBz CAR T cells exhibited the greatest expansion after secondary stimulation, followed by 19PBBz and 19TBBz CAR T cells, which showed similarly improved expansion compared with 19GBBz CAR T cells, respectively (**Fig. 4a**). Interestingly, the proliferative advantage of the 19PTBBz cells compared to 19PBBz was abrogated in the presence of CD226 blocking antibody (**Fig. 4b**), suggesting that the enhanced proliferative capacity of 19PTBBz CAR T cells compared with 19PBBz is mainly attributable to CD226 costimulatory signaling. To further confirm the functional importance of CD226 signaling, we knocked out the *CD226* gene using CRISPR/Cas9, which yielded a mixture of CD226^+^ and CD226^-^ 19PTBBz CAR T cells (**Supplementary Fig. 12a and b**). Indeed, after 2^nd^ stimulation of the CAR T mixture, intracellular immunostaining analysis revealed a reduced production of IL-2 in the CD226^-^ population compared with the CD226^+^ population (**Supplementary Fig. 12c and d**). Notably, we found that activated CAR T cells also express high levels of CD112 and CD155 (**Supplementary Fig. 13**), the ligands for CD226 and TIGIT, which can potentially contribute to costimulatory CD226 signaling *in cis* or *in trans* between adjacent CAR T cells^48,49^.

Several studies have demonstrated that a less-differentiated status (e.g., stem cell memory and central memory) in memory T cells is positively correlated with the efficacy of CAR T therapy^50,51^. Thus, we analyzed the composition of memory subsets of each CAR T construct based on the expression of C-C motif chemokine receptor 7 (CCR7) and CD45RO. Although a majority of freshly prepared CAR T cells have a central memory (T_CM_) phenotype, both CD4^+^ and CD8^+^ 19GBBz CAR T cells dramatically differentiated into the effector memory (T_EM_) phenotype upon repeated *in vitro* stimulation **(Supplementary Fig. 14**). Interestingly, however, the degree of differentiation was lower in the two TIGIT-downregulated CAR T cells, 19PTBBz and 19TBBz, compared with 19GBBz and 19PBBz, resulting in a higher T_CM_/T_EM_ ratio in both CD4^+^ and CD8^+^ populations (**Fig. 4c and Supplementary Fig. 15**).

We next compared the cytotoxic activity of each CAR T construct against Nalm-6-PD-L1 cells at various effector-to-target (E:T) ratios. In contrast to the differentiation pattern observed above, 19PTBBz and 19PBBz demonstrated similarly higher killing efficiency compared with 19TBBz and 19GBBz at a low E:T ratio (0.1:1), suggesting that PD-1 signaling, but not TIGIT, has a major impact on the short-term cytotoxicity of CAR T cells (**Fig. 4d**). Overall, the results from our *in vitro* comparisons suggest that inhibitory PD-1 and TIGIT signaling have non-redundant effects on the differentiation, proliferation, and effector functions of CAR T cells; thus, simultaneous downregulation of these two ICRs may synergistically benefit their antitumor activity.

### Downregulation of PD-1 and TIGIT distinctly reprograms the transcriptomic profiles of CAR T cells

To better understand the functional characteristics of ICR downregulation, we performed bulk RNA sequencing (RNA-seq) on CAR T cells generated from two independent donors. First, we analyzed 19GBBz CAR T cells harvested after repeated stimulation and found significant changes in their transcriptomic profiles (**Supplementary Fig. 16a-c**). Specifically, the expression of genes encoding inhibitory receptors *(PD-1, TIGIT, LAG-3, TIM-3,* and *CD244),* exhaustion-related transcription factors (PRDM1, TOX2, EOMES, and EGR2/3), and chemokines *(XCL1, CCL3,* and *CLL4)* was significantly increased, whereas the expression of genes encoding naïve/central memory-associated markers and transcription factors *(BCL6, IL7R, TCF7, LEF1, SELL, CD27,* and *CCR7*), alongside those encoding proteins associated with glucose metabolism *(HK2, PFKFPB4, PDK1, GYS1,* and *GBE1)* was decreased (**Supplementary Fig. 16d**). A gene set enrichment analysis (GSEA) also revealed that CAR T cells exhibited a more differentiated and exhausted state after repeated stimulation^52,53^ (**Supplementary Fig. 16e**).

Next, we compared the gene expression profiles of 19GBBz, 19PBBz, 19TBBz, and 19PTBBz CAR T cells after 2^nd^ stimulation with Nalm-6-PD-L1 cells. A Pearson’s correlation analysis revealed significant similarity between the two TIGIT-downregulated CAR T cells 19TBBz and 19PTBBz relative to 19GBBz and 19PBBz – indicating that the high degree of transcriptomic reprogramming of CAR T cells is mainly regulated by TIGIT rather than PD-1 downregulation (**Fig. 5a**). We then analyzed the hierarchically clustered heatmap of differentially expressed genes in each group of CAR T cells (**Fig. 5b**). Notably, the PD-1– downregulated CAR T cells, 19PBBz and 19PTBBz, showed similar increases in the number of transcripts for effector and proliferation-related molecules *(IL2, MKI67, FASLG,* and *TNFSF10).* In contrast, transcriptional features related to the differentiation and exhaustion status of T cells were largely shared between 19TBBz and 19PTBBz CAR T cells but were clearly distinct from those of 19GBBz and 19PBBz CAR T cells. In general, 19TBBz and 19PTBBz CAR T cells exhibited decreased expression of inhibitory receptor *(LAG3* and *CD244)* and chemokine genes *(XCL1, XCL2,* and *CCL3*) and higher expression of naïve/central memory-phenotype *(IL7R, BCL6,* and *CD27)* and active glucose metabolism genes *(HK2, PFKFB4,* and *PDK1),* indicating a less-differentiated/exhausted state (**Fig. 5c)**. GSEA also revealed the downregulation of exhaustion-related genes and the upregulation of naïve/memory-related genes in 19PTBBz CAR T cells compared with 19PBBz (**Fig. 5d**). These RNA-seq results are consistent with our *in vitro* functional studies and further support our hypothesis that the downregulation of PD-1 enhances short-term effector function in 19PTBBz CAR T cells, while the downregulation of TIGIT is primarily responsible for maintaining a less differentiated/exhausted state.

### *In vivo* efficacy of patient-derived, clinical-grade PD-1/TIGIT–downregulated CAR T cells

To assess the feasibility of the clinical translation of PD-1/TIGIT dual-downregulated CD19-targeting CAR T cells, we established a large-scale CAR T manufacturing protocol using a semi-automated closed system (**Supplementary Fig. 17a**). In this protocol, we used the modified lentiviral vectors 19PBBz-nt and 19PTBBz-nt in which the ΔLNGFR tag, which was inserted for purification of CAR T, was removed from the original constructs. For comparison, we also generated 19BBz-nt without an shRNA cassette (**Supplementary Fig. 17b**). Consistent with the results obtained above, healthy donor-derived 19PTBBz-nt CAR T cells outperformed 19BBz-nt and 19PBBz-nt in our Nalm-6-PD-L1 leukemia model (**Supplementary Fig. 17c**). Next, following the same protocol, we manufactured 19BBz-nt and 19PTBBz-nt CAR T from diffuse large B-cell lymphoma (DLBCL) patient-derived T cells (**Supplementary Fig. 18**). After 5 days of *ex vivo* culture, CAR T cells were successfully manufactured with a similar transduction efficiency (26.8% ± 11.44% for 19PTBBz-nt and 32.3% ± 11.07% for 19BBz-nt, respectively, **Supplementary Fig. 18a**), CD4/CD8 ratio, and differentiation profiles (**Supplementary Fig. 18b and c**). Cell growth was also comparable as determined by absolute cell count and glucose consumption (**Supplementary Fig. 18d and e**). However, robust downregulation of PD-1 and TIGIT was observed only in the CAR^+^ population of the 19PTBBz-nt group (**Supplementary Fig. 18f**). Both groups showed persistent Nalm-6-PD-L1 tumor clearance in all mice treated with a dose of 1×10^6^ CAR T cells. However, mice treated with 19BBz-nt experienced tumor relapse at a dose of 0.5×10^6^ CAR T cells and almost lost tumor control at a dose 0.25×10^6^. In contrast, 19PTBBz-nt CAR T cells effectively suppressed tumor relapse even at a dose of 0.25×10^6^ CAR T cells (**Fig. 6a and b**). Mouse survival was also significantly improved by treatment with 19PTBBz-nt CAR T cells compared to 19BBz-nt at a dose of 0.25×10^6^ CAR T cells (**Fig. 6c**), a finding that may be explained by the difference in the persistence CAR T cells *in vivo* at a dose of 0.5×10^6^ (**Fig. 6d**). Lastly, we evaluated the *in vivo* efficacy of 19PTBBz-nt and 19BBz-nt in a subcutaneous Raji-PD-L1 lymphoma model and confirmed the improved antitumor activity of 19PTBBz-nt CAR T cells over 19BBz-nt across different doses (4×10^6^, 2×10^6^, and 1×10^6^ CAR T cells, **Fig. 6f and g**).

Collectively, these results show that our cell-intrinsic PD-1/TIGIT dual-downregulation strategy, which is readily applicable to a clinical-grade CAR T manufacturing process, may provide an effective approach for enhancing the *in vivo* efficacy of CAR T cells by averting immune checkpoint-mediated dysfunction.

## Discussion

Since the first historic approval of CD19-targeting CAR T cell therapy for B-cell malignancies in 2017, several lines of preclinical and clinical evidence have strongly suggested that the efficacy of CAR T cells should be further improved to benefit a broader range of cancer patients, including those with solid tumors. Because inhibitory checkpoint signaling in CAR T cell is considered one of the reasons for this suboptimal efficacy, we applied an shRNA-based gene-silencing approach to viral vector design to achieve sustained downregulation of checkpoint receptors while minimizing systemic toxicity. An alternative approach for achieving the same goal is applying genome-editing tools such as CRISPR/Cas9 for complete gene loss of function, which is widely used in the development of T cell therapeutics, including CAR T. Compared with shRNA-mediated knockdown, CRISPR/Cas9 induces a permanent knockout of target genes in the edited cell population. This would be preferable in cases where the complete loss of function of the target gene is unquestionably necessary^54^. However, in a mouse model of chronic viral infection, the genetic absence of PD-1 in antigen-specific CD8 T cells significantly reduces long-term T cell survival, likely owing to chronic overstimulation, despite their enhanced cytotoxicity and proliferation during the acute phase of infection^55^. Similarly, complete deficiency in thymocyte selection associated high mobility group box (TOX) or eomesodermin (EOMES), the key transcription factors that promote high expression of inhibitory receptors and T cell exhaustion, results in a more rapid decline of antigen-specific T cells in mouse models of chronic viral infection and cancer^56–58^. However, deletion of only one allele of TOX or EOMES was shown to rescue the persistence of antigen-specific T cells, resulting in superior disease control compared with wild-type T cells^57,58^. These results strongly suggest that the partial inhibition of exhaustion-associated factors in engineered T cells may not only be sufficient, but also beneficial for induction of optimal therapeutic outcomes. In addition, our approach to integrating shRNA cassettes into lentiviral vectors is readily applicable to currently established protocols for commercial CAR-T manufacturing; moreover, its manufacturing costs and risk of failure could be relatively lower compared with CRISPR-based knockout, which requires additional steps and reagents.

We showed that cell-intrinsic PD-1 downregulation endowed 41BB-based CD19-targeting CAR T cells with resistance to the inhibitory PD-1/PDL-1 signaling axis, and the resulting 41BB-based CAR-T cells exerted more persistent *in vitro* and *in vivo* activity compared with their CD28-based counterpart. This is consistent with results of a previous comparison of conventional 41BB- and CD28-based CAR T cells ^59^. Next, to assess the effects of dual ICR downregulation, we compared the combination of PD-1 with TIM-3, LAG-3, TIGIT, and CTLA-4, which have been demonstrated to have synergistic effects in several preclinical models and/or clinical trials as antibody-based blockade targets when combined with anti-PD-1 antibodies^19,22,23,60–62^. However, among the four different combinations, only PD-1/TIGIT enhanced the antitumor activity of CD19-targeting CAR T cells compared with single downregulation of PD-1. We cannot currently rule out the possibility that these results are attributable to limitations in the mouse xenograft model used. Although we confirmed that our Nalm-6-GL-PD-L1 cell line expresses galectin-9, HLA-DR, and CD112, which are the respective ligands for TIM-3, LAG-3, and TIGIT, B7, the ligand for CTLA-4, was not detected. Furthermore, the degree of cross-species reactivity between checkpoint receptors in human CAR T cells and their ligands expressed in mouse tissue may be variable depending on the receptor type, thereby biasing the experimental results.

Nonetheless, it is also plausible that our results reflect a biological feature of CAR T cells that distinguishes them from endogenous tumor-reactive T cells. For example, anti-CTLA-4 antibodies are known to exert their effects primarily by rescuing the costimulatory CD28/B7 axis between naïve T cells and activated dendritic cells (DCs) within the priming site^15^, which may not be relevant for CAR T cells with a 41BB costimulatory domain that has undergone *ex vivo* stimulation by anti-CD3 and anti-CD28 antibodies. Similarly, HLA-DR, the ligand for LAG-3, is also highly expressed on antigen-presenting cells, and it has been shown that the therapeutic effect of LAG-3 blockade is mediated by enhancing T cell activation by these cells^61^, which may also not be relevant for CAR T cells.

Our results may further be explained by mechanistic differences between our cell-intrinsic downregulation approach and systemic antibody treatment. A recent clinical study showed that combination therapy using anti-PD-1 and anti-CTLA-4 antibodies induced different therapeutic effects depending on the treatment sequence^22^. This may suggest that each antibody used in the combination therapy exerts a synergistic effect by engaging distinct T cell populations undergoing different activation and differentiation stages, rather than engaging the same T cells simultaneously. Lastly, although it is well known that treatment with TIM-3–targeting antibodies enhances the effector function of T cells and induces antitumor responses, both stimulatory and inhibitory functions of TIM-3 in T cells have been reported^63,64^. These contradictory results, together with the fact that there are no known inhibitory signaling motifs in the cytoplasmic domain of TIM-3, raise the possibility that TIM-3–targeting antibodies may in fact act as agonists^65^.

*In vitro* functional assays and RNA-seq analysis revealed that downregulation of PD-1 enhances effector function whereas downregulation of TIGIT is primarily responsible for the acquisition of a less-differentiated/exhausted phenotype. There have been several reports that inhibition of PD-1 signaling enhances the effector function of T cells^15,25,27,33,66,67^. Interestingly, in our experimental conditions, PD-1 downregulation did not affect the transcription of the soluble effectors interferon-γ, granzyme B, and perforin 1, but did promote the transcription of the membrane-bound death ligands TNF superfamily member 10 (TNFSF10) and Fas ligand (FASLG). It was recently reported that genetic disruption of TNFSF10 or FASLG in CAR T cells significantly impairs cytotoxic activity and induces the progressive dysfunction of CAR T cells^68^.

In the case of TIGIT, several mechanisms have been proposed for its inhibitory effects. First, it can impair the effector functions of T and NK cells through its intracellular ITIM domain or through competition with the costimulatory receptor CD226^49,69^. It also has been reported that TIGIT regulates immune responses by modulating DCs and regulatory T cells (Tregs)^69,70^. Interestingly, in the study by Kurtulus et al., induction of TIGIT signaling by an agonistic antibody was shown to decrease the levels of the transcription factor TCF-1^70^, a decrease which, in CD8 T cells, inhibits differentiation into effector cells while promoting the development of a central memory phenotype^71^. In addition, a recent study showed that treatment with an agonistic antibody targeting glucocorticoid-induced tumor necrosis factor receptor-related protein (GITR) in combination with a PD-1-blocking antibody generated memory CD8 T cells with high proliferative capacity, resulting in a synergistic regression of tumors^72^. Notably, the authors showed that the effect was predominantly dependent on CD226 signaling, which is enhanced by both GITR and PD-1 blockade; this finding is also in line with our results and those of others highlighting the role of CD226 in TIGIT blockade^73^. Collectively, these reports alongside our data strongly suggest that one of the principal mechanisms of actions of TIGIT blockade can be the modulation of the differentiation and memory status of tumor-reactive T cells.

In recent clinical trials, combination therapy using anti-PD-L1 and anti-TIGIT blocking antibodies has shown promising results in terms of both efficacy and safety, raising expectations for this particular combination of checkpoint receptors. To the best of our knowledge, our study is the first to report that downregulation of these two ICRs in a cell-intrinsic fashion synergistically enhances the antitumor activity of CAR T cells. Our study also provides a mechanistic rationale for the synergy, laying the groundwork for more rigorous future single-cell-level analyses and confirmation through alternative blockade strategies such as knockout or antibody treatment. Moving forward, whether PD-1/TIGIT dual-downregulated CAR T cells will be able to induce a more robust response in patients who were refractory to previous CAR T therapies should be determined in a clinical setting. Furthermore, whether this combination can be applied to other engineered T cell platforms, such as TCR T therapy, as well as to CAR T cells targeting solid tumor antigens, is a subject of considerable interest that is currently under investigation in preclinical models.

## Supplementary Figure Legends

**Supplementary figure 1.**
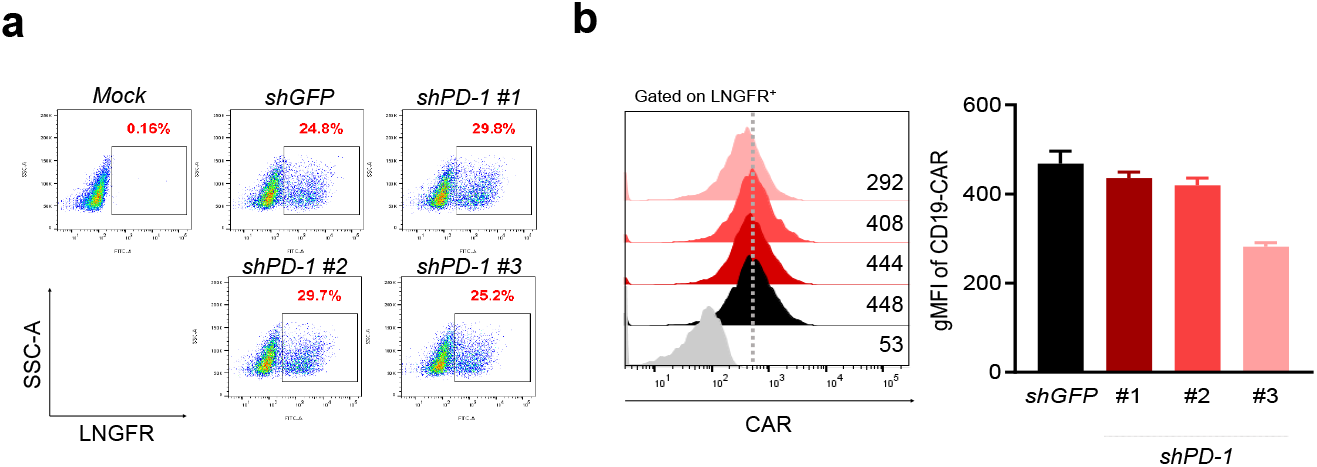
Generation of PD-1-downregulated CAR T cells. **(a)** Transduction efficiency of CAR T cells transduced with empty (mock), GFP, or PD-1 shRNA #1-3-containing lentiviruses was determined as the percentage of ΔLNGFR^-^ positive cells. **(b)** Surface CAR expression was analyzed with APC-conjugated anti-mouse Fab antibody on day 6 after the LNGFR^+^ isolation of transduced cells. Error bars represent the range from n = 2 independent donors in separate experiments.

**Supplementary figure 2.**
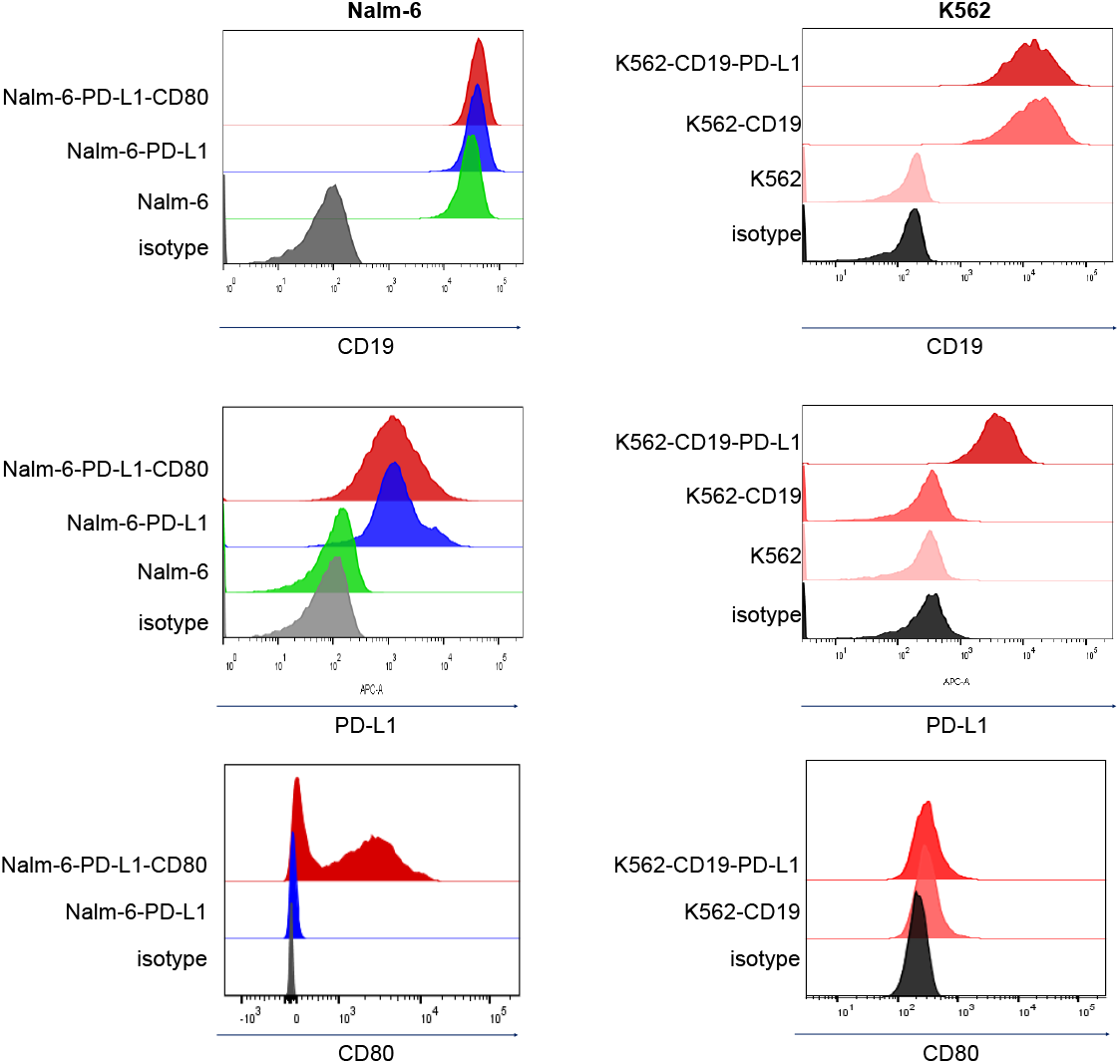
Target cell expression of CD19, PD-L1, and CD80. Surface CD19, PD-L1, and CD80 expression levels of Nalm-6-GL-PD-L1-CD80, Nalm-6-GL-PD-L1, Nalm-6, K562-CD19-PD-L1, K562-CD19, and K562 cells was determined by flow cytometry.

**Supplementary figure 3.**
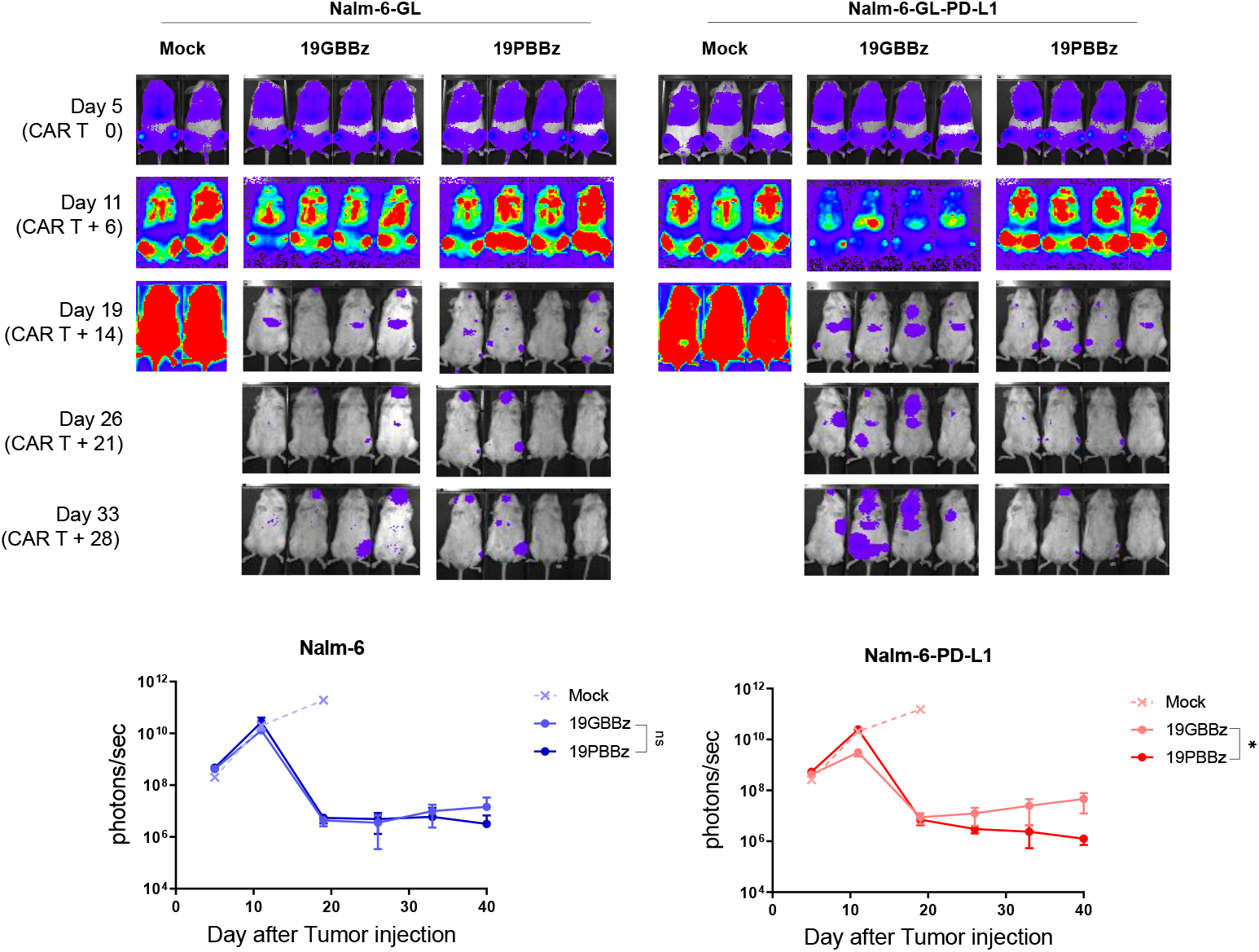
PD-1 downregulation enhances the *in vivo* antitumor activity of CAR T cells in a manner dependent on target cell expression of PD-L1. 1×10^6^ Nalm-6-GL or Nalm-6-GL-PD-L1 leukemia cells were injected intravenously into NSG mice. 5 days later, 1×10^6^ CAR T cells were injected intravenously. Tumor burden was monitored based on the bioluminescence intensity from the IVIS imaging system. Data are from n = 2 mock-, n = 4 19GBBz-, and n = 4 19PBBz-treated Nalm-6-GL bearing mice and n = 3 mock-, n = 4 19GBBz-, and n = 4 19PBBz-treated Nalm-6-GL-PD-L1 bearing mice. Statistical analysis was done by unpaired two-tailed t-test on the results from day 40.

**Supplementary figure 4.**
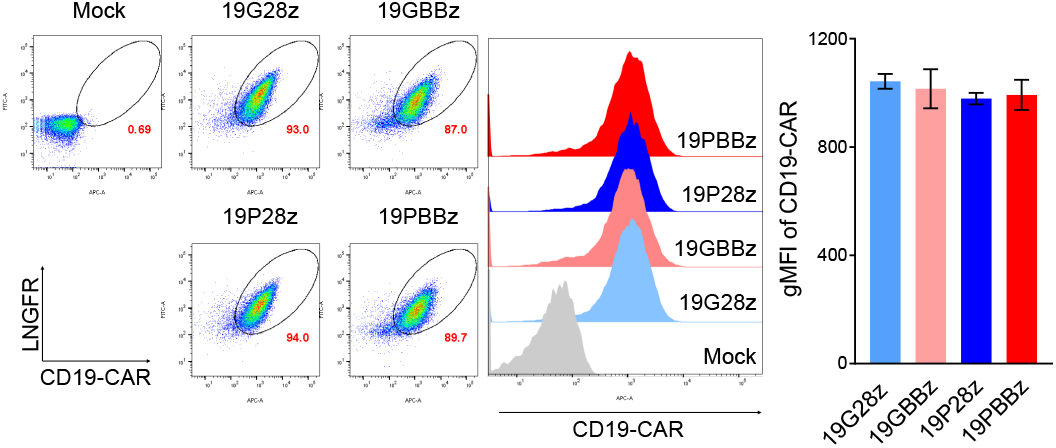
CD19 specific CAR expression level of 19G28z, 19GBBz, 19P28z, and 19PBBz cells. CAR and ΔLNGFR expression level of 19G28z, 19GBBz, 19P28z, and 19PBBz cells on day 6 following LNGFR^+^ isolation. Bars represent the ranges from CAR T cells generated from two donors.

**Supplementary figure 5.**
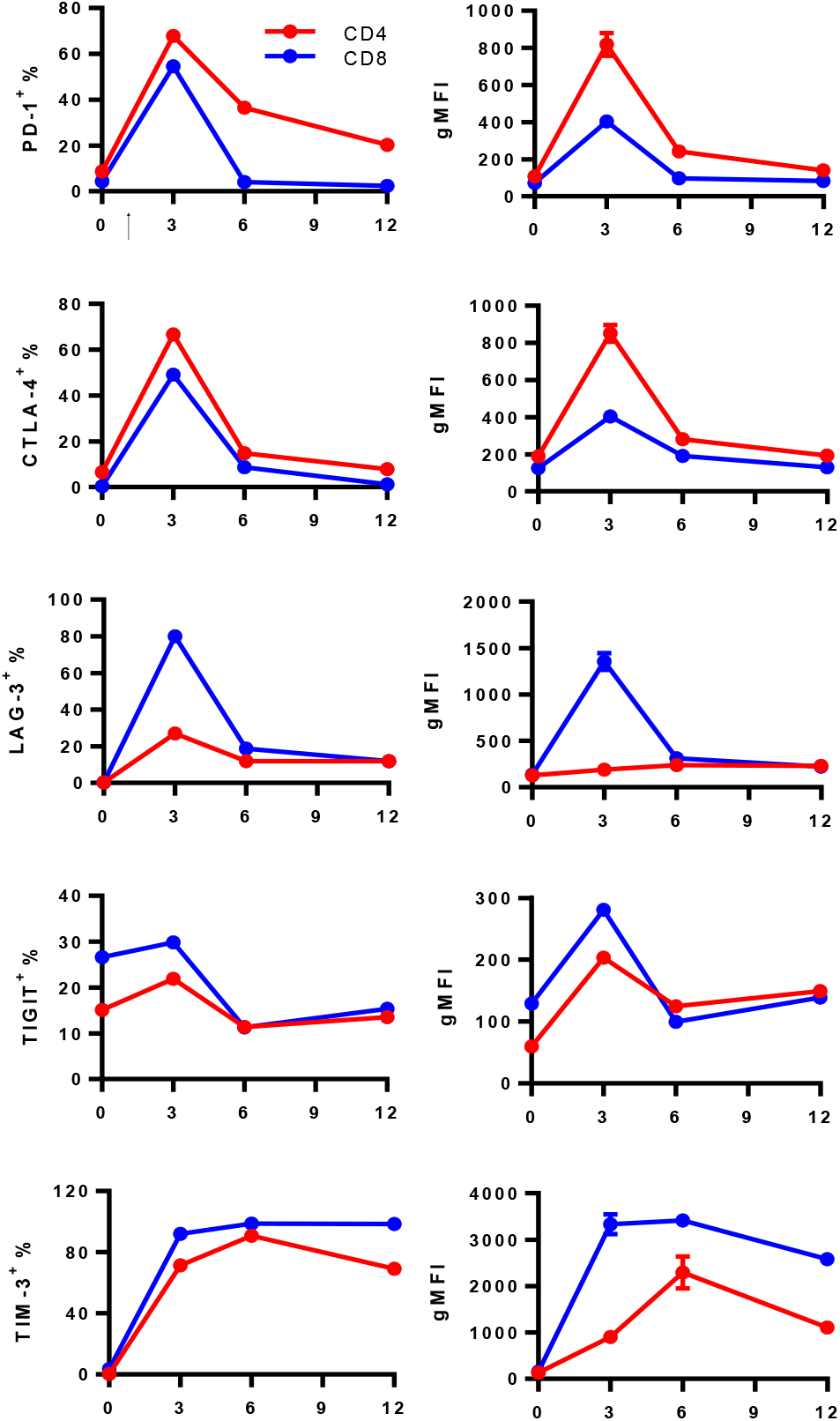
Kinetics of PD-1, CTLA-4, LAG-3, TIGIT, and TIM-3 expression in activated CD4 or CD8 T cells. PBMCs were stimulated with anti-CD3/CD28 beads. PD-1, LAG-3, TIGIT, and TIM-3 expression levels of CD4^+^ and CD8^+^ T cells were determined by flow cytometry on days 0, 3, 6, and 12 after stimulation. The CTLA-4 expression level was evaluated through intracellular staining.

**Supplementary figure 6.**
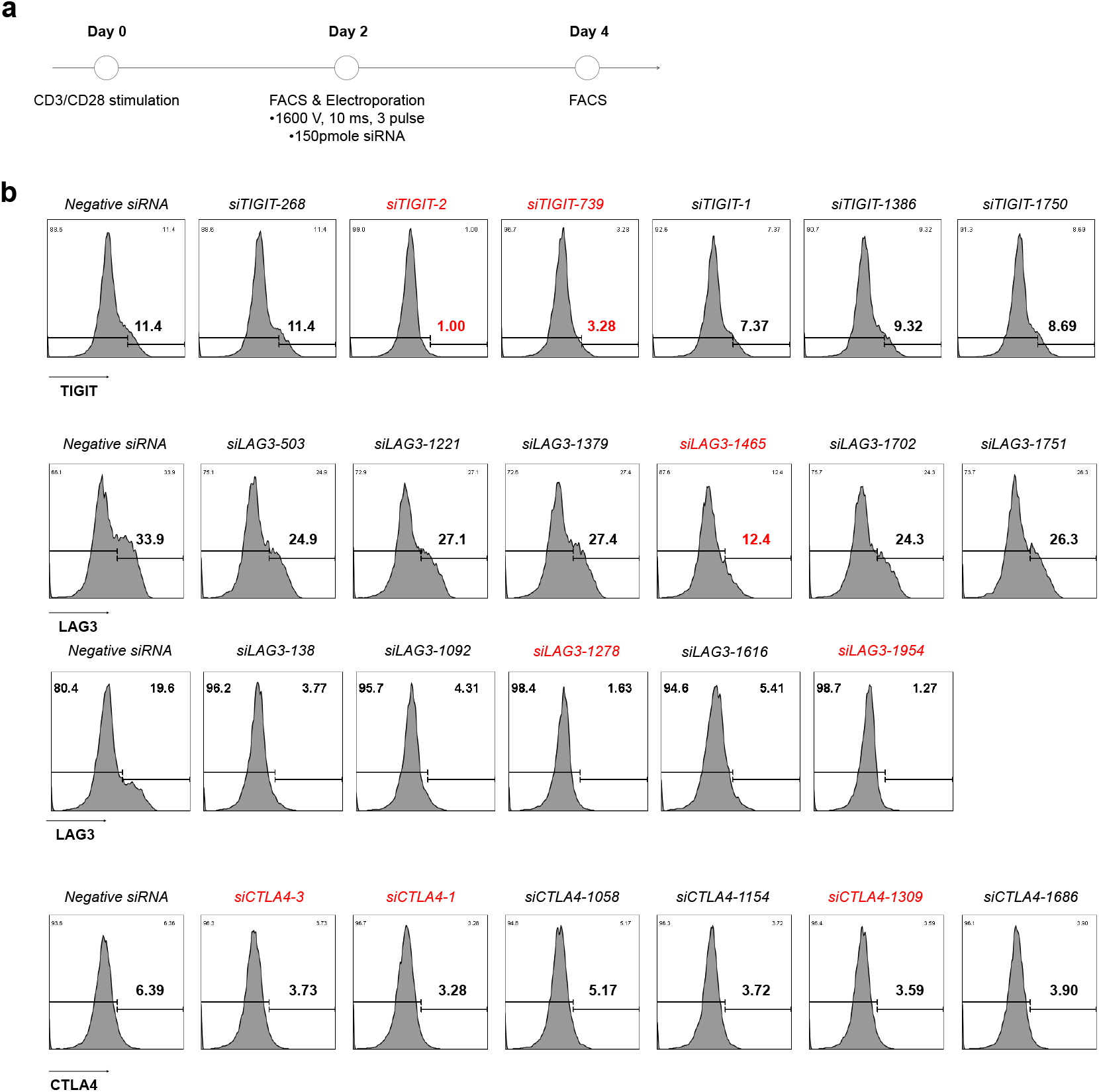
Selection of 21-mer siRNA candidates targeting TIGIT, LAG-3, and CTLA-4. **(a)** Schematic representation of siRNA screening and selection. 150 pmol siRNA candidates were electroporated (Neon electroporation system, 1600V, 10ms, 3 pulses) into T cells 2 days after stimulation. The expression level of inhibitory receptors was evaluated by flow cytometry on day 4. siRNA targeting GFP was used as a negative control. **(b)** Representative FACS plots of TIGIT, LAG-3, and CTLA-4 expression in T cells electroporated with each siRNA.

**Supplementary figure 7.**
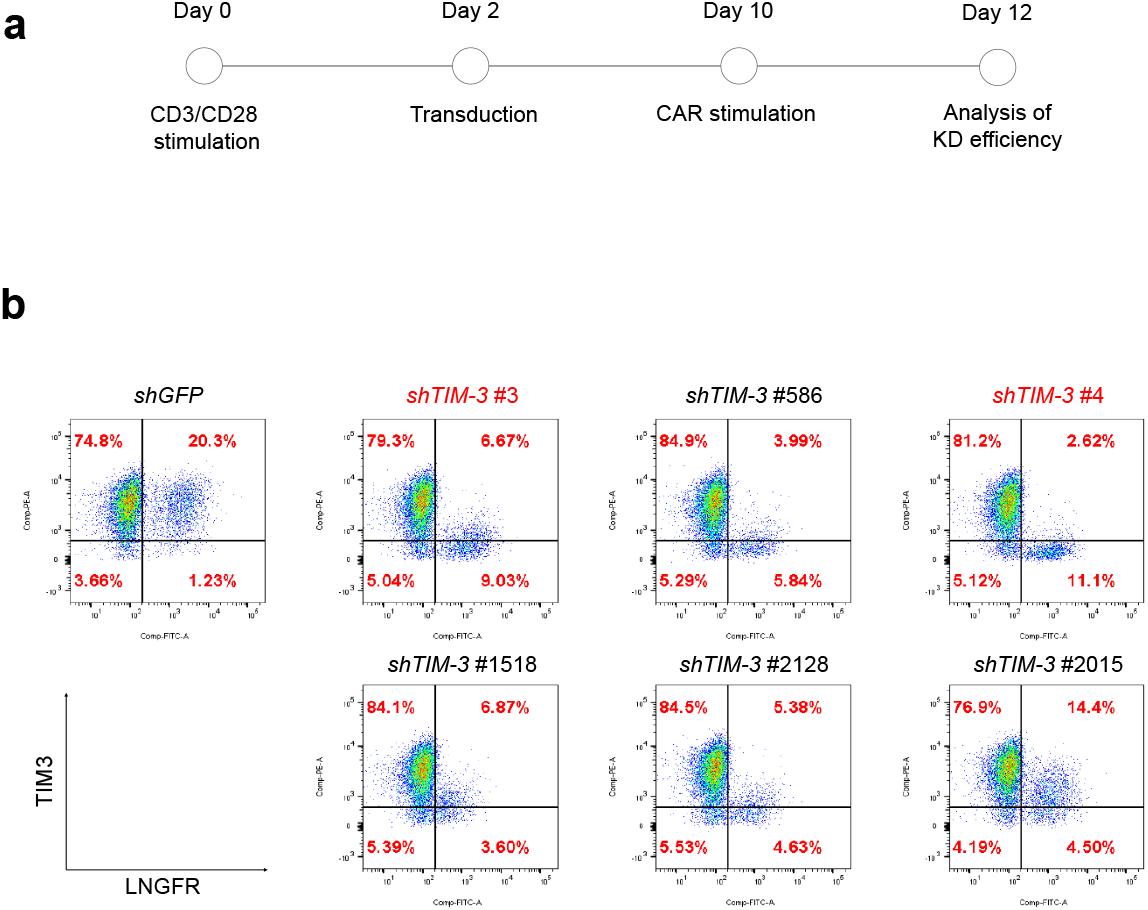
Selection of shRNA candidates targeting TIM-3. **(a)** Schematic representation of the generation and analysis of *shTIM-3-CD19* CAR constructs as described in Supplementary Figure 1. CD19-specific CAR T cells were stimulated with γ-irradiated K562-CD19 cells for 2 days and analyzed for the expression of TIM-3 by flow cytometry, **(b)** Representative FACS plots of TIM-3 expression in ΔLNGFR^+^ CAR T cells expressing each TIM-3 targeting shRNA. shRNA targeting GFP *(shGFP)* was used as a negative control.

**Supplementary figure 8.**
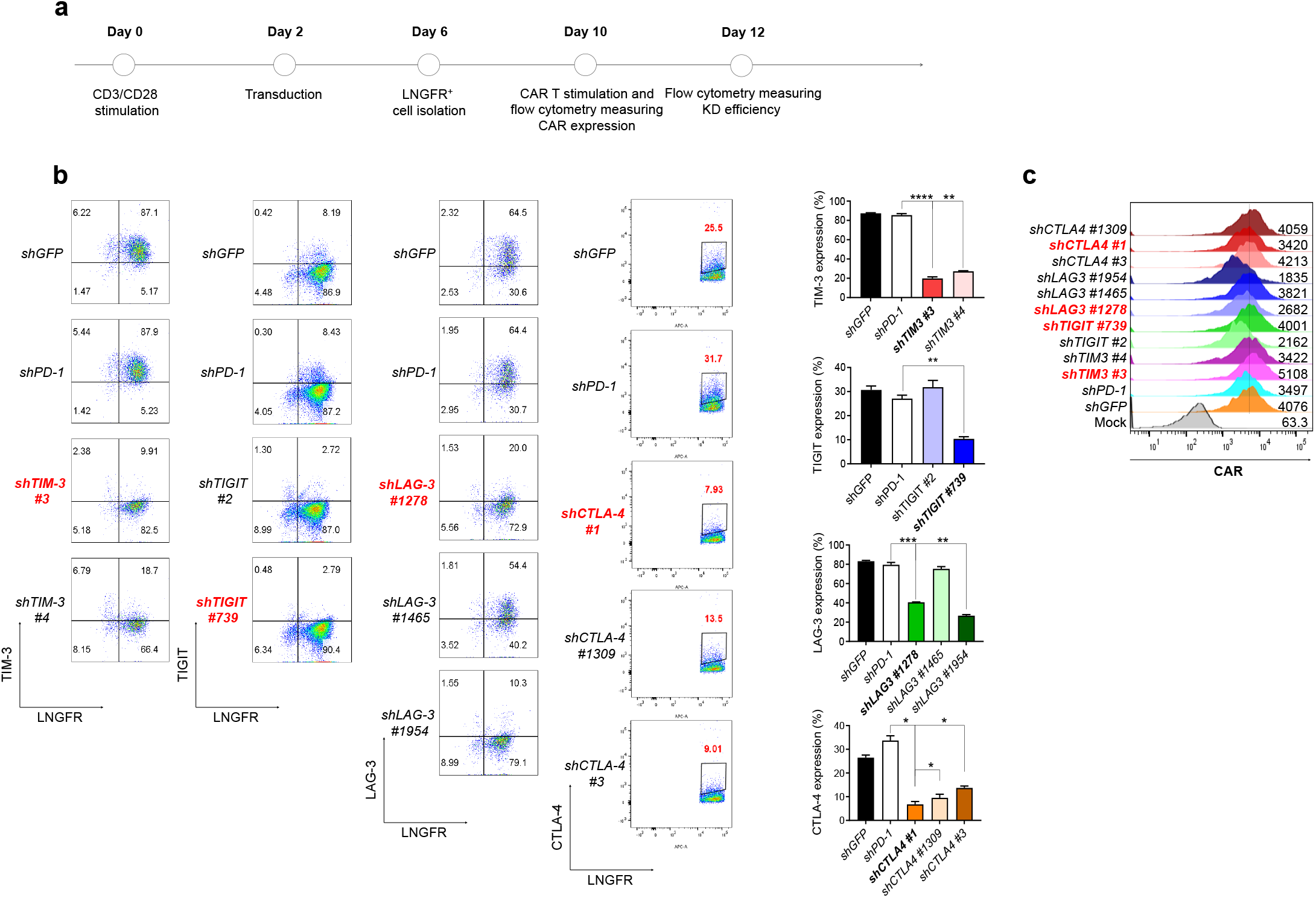
Knockdown efficiency of ICRs and CAR expression in CD19 specific CAR T cells. **(a)** Schematic representation of the generation of CD19-specific CAR T cells with selected shRNA sequences targeting TIM-3, TIGIT, LAG-3, and CTLA-4. ΔLNGFR^+^ T cells were stimulated with γ-irradiated K562-CD19 cells on day 10 and knockdown efficiency was measured on day 12. **(b)** The expression level of inhibitory receptors in CAR T cells expressing the indicated shRNA cassetttes was evaluated by flow cytometry on day 12. **(c)** CAR expression levels in CAR T cells expressing the indicated shRNA cassettes were determined on day 10 by flow cytometry. Data are the pooled mean ± SD from two indepdendent experiments performed in duplicates. Statistical analysis was done by One-Way ANOVA. * p < 0.05, ** p < 0.01, *** p < 0.001, **** p < 0.0001.

**Supplementary figure 9.**
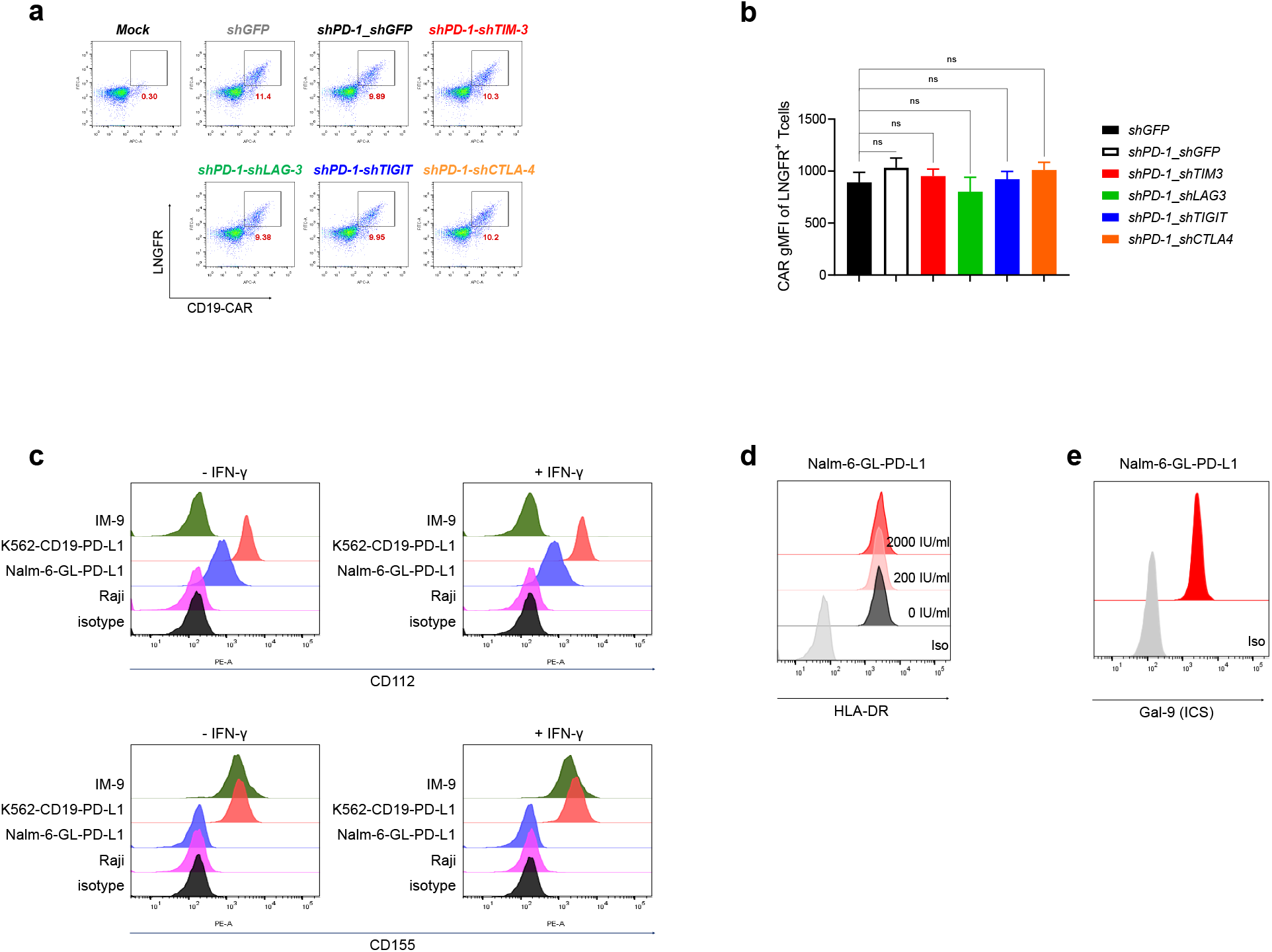
Generation of CAR T cells with dual downregulation of inhibitory receptors. **(a)** The transduction efficiency of dual shRNA constructs was determined by measuring CD19 CAR ΔLNGFR expression by FACS on day 4 after transduction. **(b)** The gMFI of CD19-CAR in ΔLNGFR^+^ T cells on day 10 after transduction. Data are from the mean ± SD from three donors. **(c)** CD112 or CD155 expression level in Raji, Nalm-6-GL-PD-L1, K562-CD19-PD-L1, and IM-9 cells with or without IFN-γ treatment for 24 h. CD112 or CD115 expression **(d)** HLA-DR expression level in Nalm-6-GL-PD-L1 cells with or without IFN-γ treatment at the indicated doses for 24 hours. **(e)** The intracellular expression level of galectin-9 in Nalm-6-GL-PD-L1 cells. Statistical analysis was done by One-Way ANOVA. ns, not significant.

**Supplementary figure 10.**
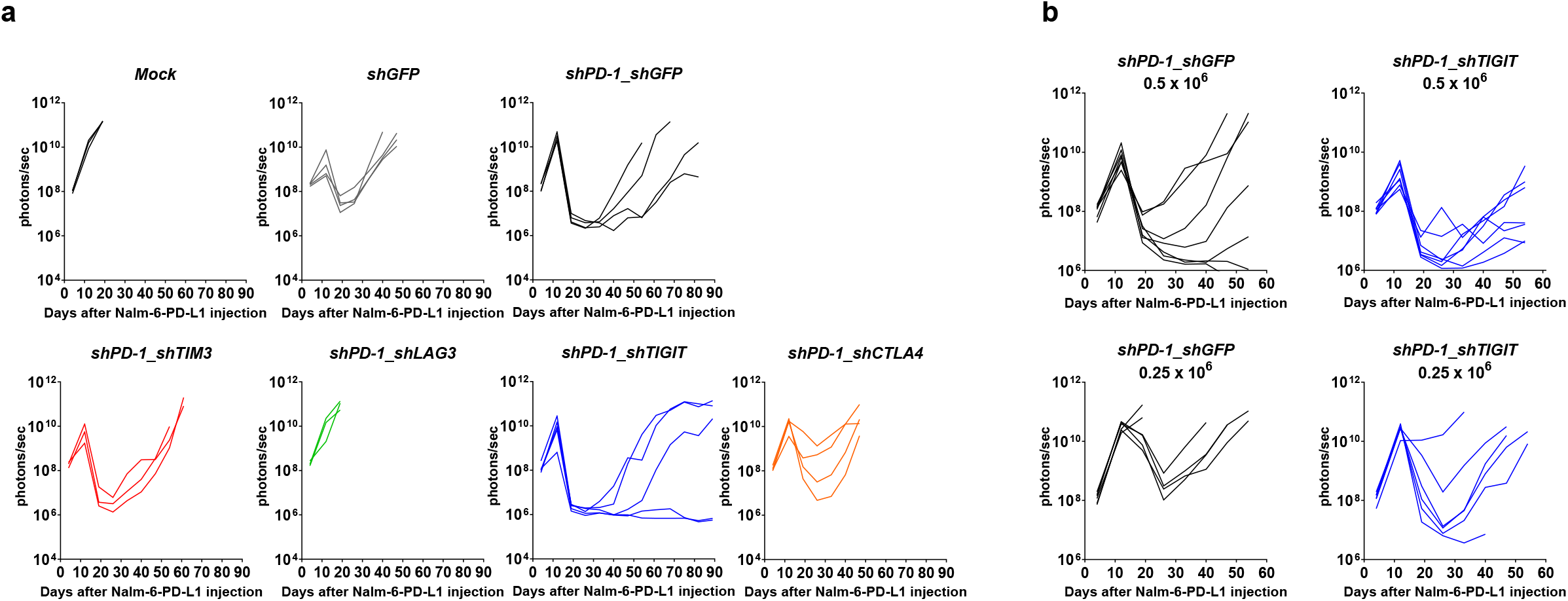
Analysis of tumor burden performed in Figure 3. **(a)** NSG mice were injected intravenously with 1×10^6^ Nalm-6-GL-PD-L1 leukemia cells. 5 days later, 1×10^6^ CAR T cells with dual downregulation *(shGFP, shPD-1_shGFP, shPD-1_shTIM-3, shPD-1_shTIGIT, shPD-1_shLAG-3, shPD-1_shCTLA-4)* were injected intravenously. Tumor burden was monitored using the bioluminescence IVIS imaging system for 89 days following tumor engraftment. **(b)** Nalm-6-GL-PD-L1 bearing mice were treated with 0.5 × 10^6^ or 0.25 × 10^6^ CAR T cells with PD-1 or PD-1/TIGIT downregulation. Tumor burden was monitored based on the bioluminescence intensity from the IVIS imaging system for 54 days following tumor engraftment.

**Supplementary figure 11.**
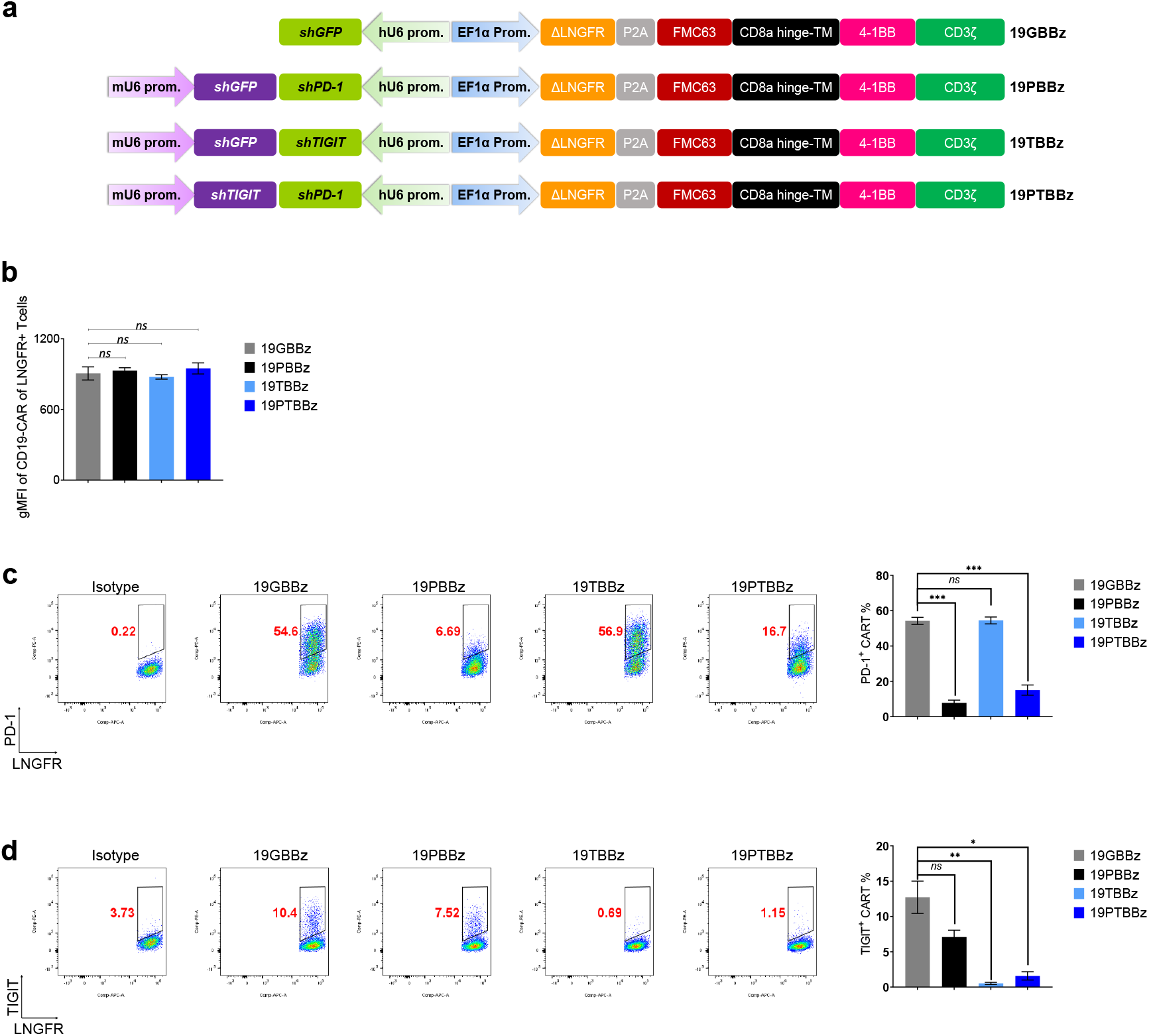
Generation and characterization of 19GBBz, 19PBBz, 19TBBz, and 19PTBBz cells. **(a)** Schematic illustration of the vector constructs used to generate 19GBBz, 19PBBz, 19TBBz, and 19PTBBz CAR T cells. **(b)** Surface expression level of CD19 CAR was evaluated by flow cytometry on day 6 after isolation of transduced cells. Data are the pooled mean ± SD from two independent experiments performed in duplicates. **(c)** PD-1 and **(d)** TIGIT expression levels of 19GBBz, 19PBBz, 19TBBz, and 19PTBBz cells stimulated with γ-irradiated K562-CD19 cells for 2 days. Data are the pooled mean ± SD from two indepdendent experiments performed in duplicates. Statistical analysis was done by One-Way ANOVA. * p < 0.05, ** p < 0.01, *** p < 0.001, ns = not significant.

**Supplementary figure 12.**
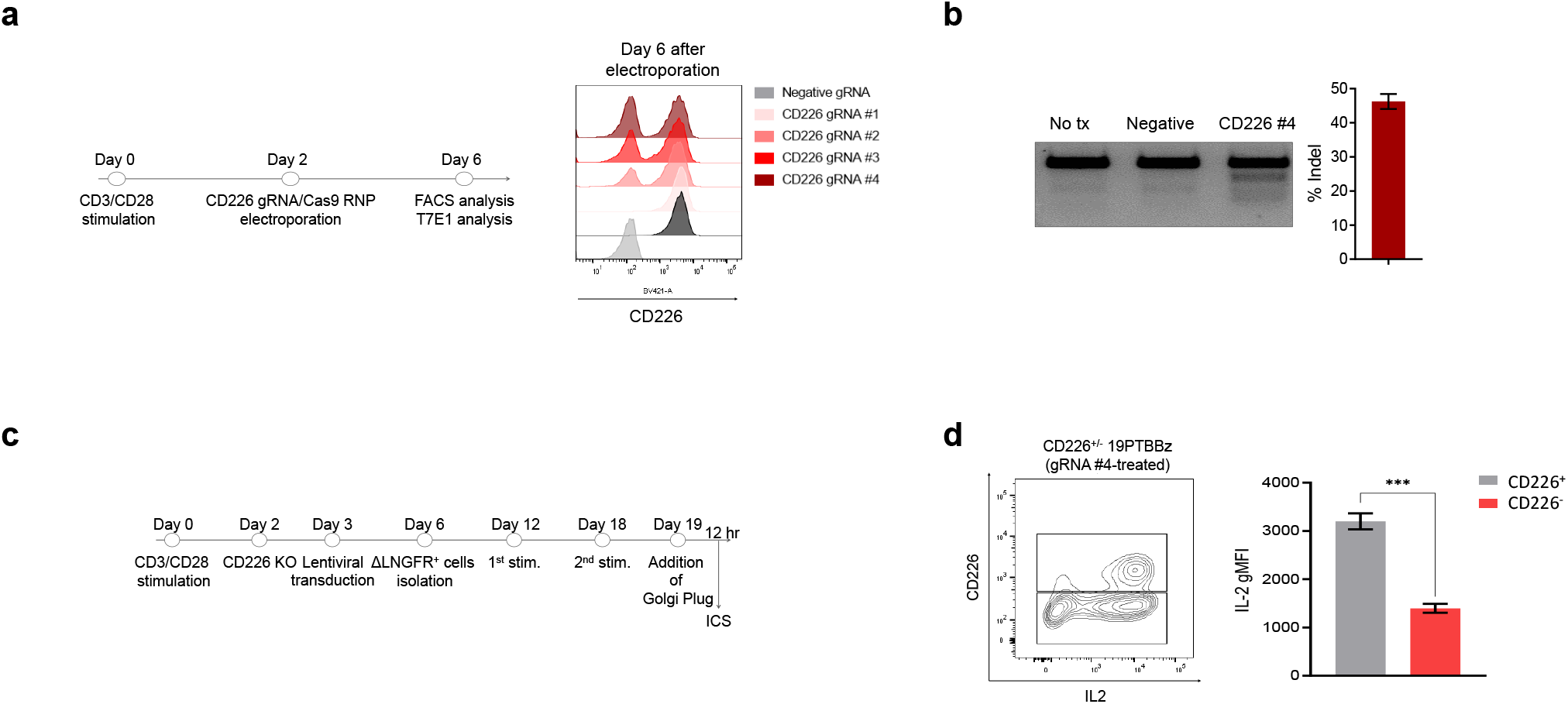
CD226 knockout and its effect on the production of IL-2 in 19PTBBz cells. **(a)** Schematic representation of CD226 knockout by CRISPR/Cas9 and the flow cytometric evaluation of the expression level of CD226 following knockout by 4 different sgRNA candidates. sgRNA targeting CAG (CMV-IE, chicken actin, rabbit beta globin) was used as a negative control. **(b)** The knockout efficiency of gRNA #4 targeting CD226 was estimated by T7 endonuclease I assay. **(c)** Schematic representation of the generation of CD226 KO CD19 specific CAR T cells and the measurement of intracellular IL-2 by flow cytometry, **(c)** Intracellular IL-2 expression levels in CD226^+^ and CD226^-^ populations measured by flow cytometry. Data are the mean ± SD from one experiment performed in triplicates.

**Supplementary figure 13.**
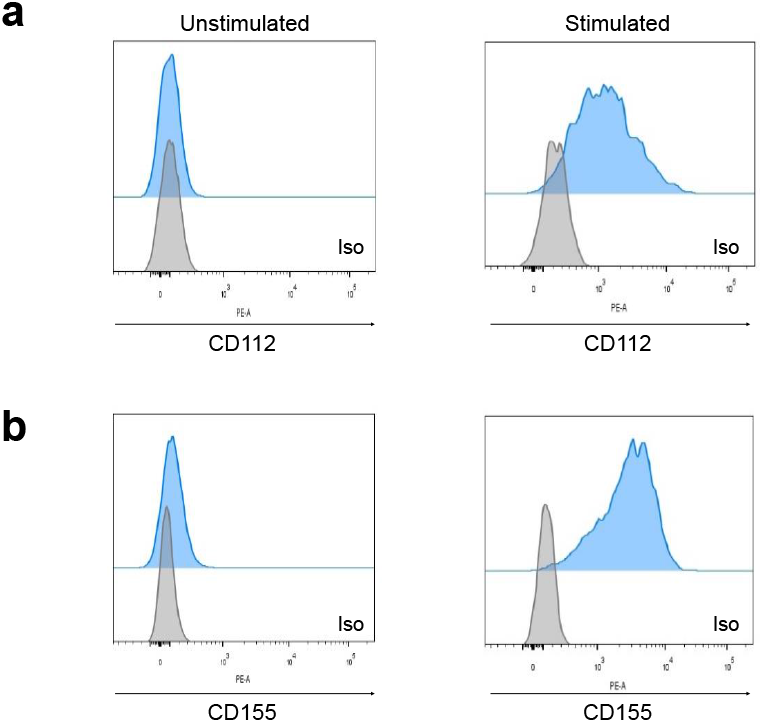
CD112 and CD155 expression on CAR T cells. Expression levels of CD112 and CD155 in 19GBBz cells with or without CD3/CD28 stimulation for 2 days measured by flow cytometry.

**Supplementary figure 14.**
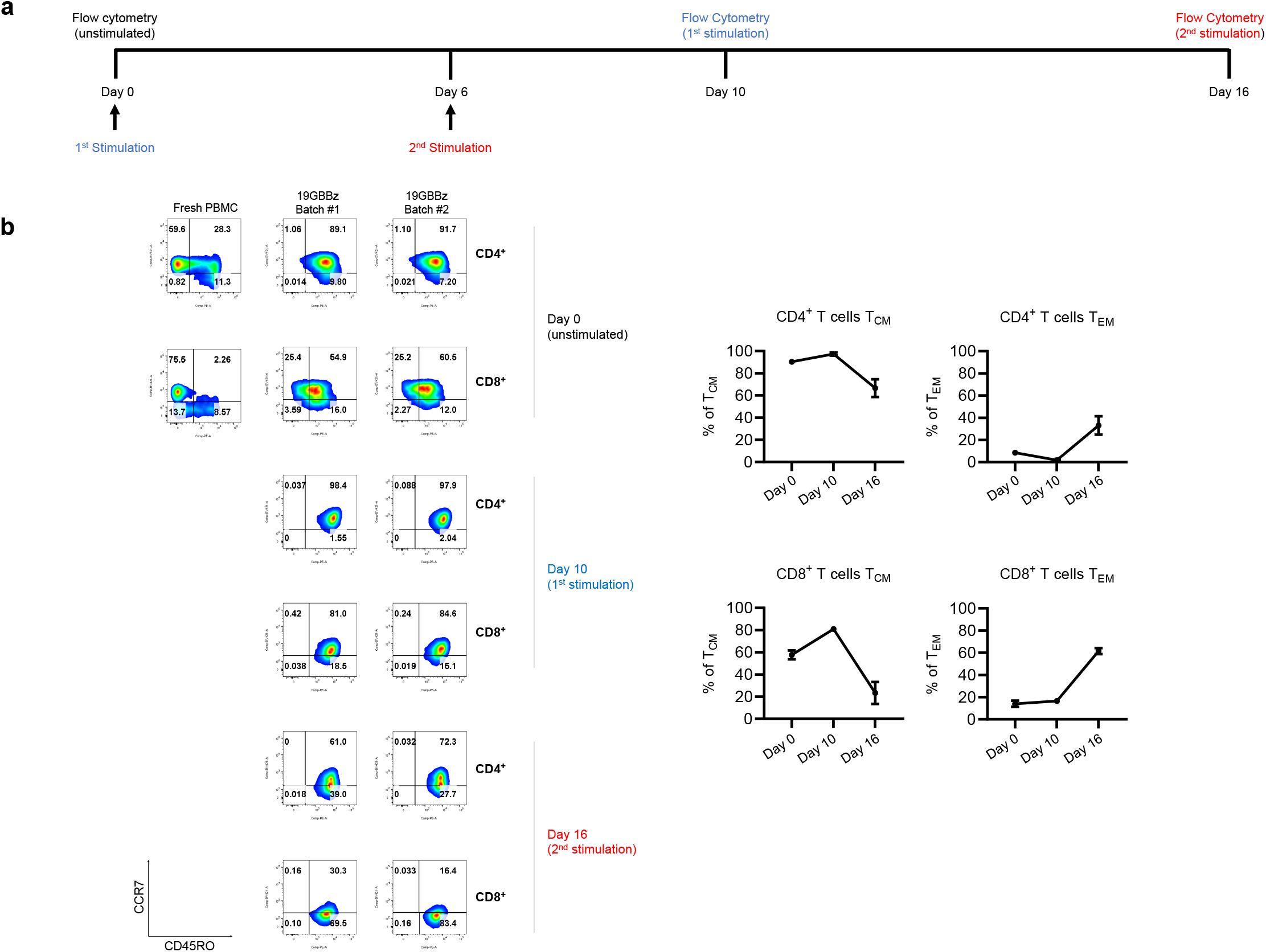
Differentiation state of 19GBBz cells upon repeated stimulation *in vitro*. 19GBBz cells were stimulated with γ-irradiated Nalm-6-GL-PD-L1-CD80 cells every six days. The differentiation state of CD4^+^ and CD8^+^ 19GBBz CAR T cells was determined by flow cytometry for CCR7 and CD45RO expression on day 10 after each stimulation. Data are the mean ± range from two donors. T_CM_ = CD45RO^+^CCR7^+^ cells, T_EM_ = CD45RO^+^CCR7^-^ cells.

**Supplementary figure 15.**
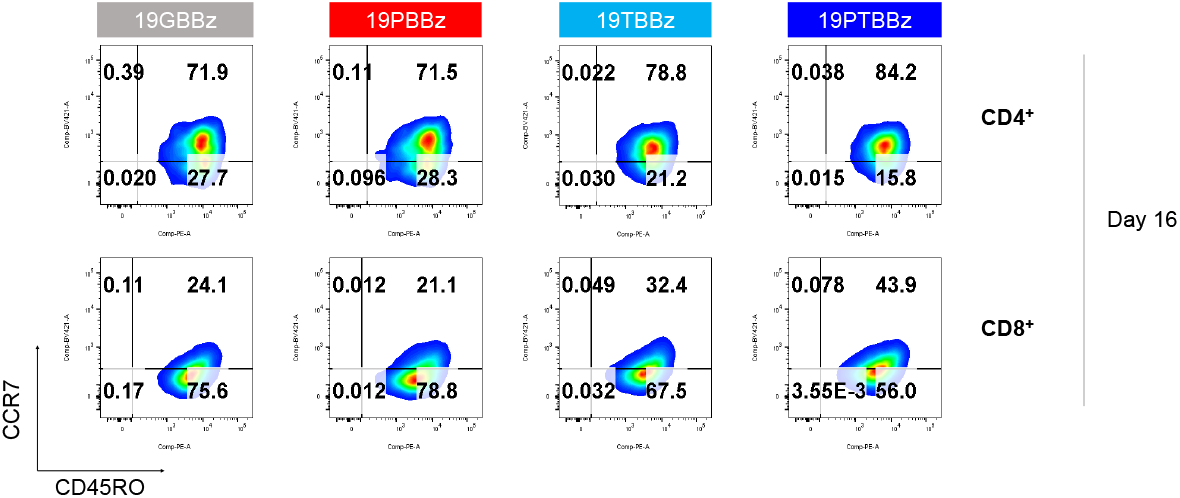
CD45RO and CCR7 expression in 19GBBz, 19PBBz, 19TBBz, and 19PTBBz cells. Representative flow cytometry plot of CD45RO and CCR7 expression in 19GBBz, 19PBBz, 19TBBz, and 19PTBBz cells on day 16 (10 days after 2^nd^ stimulation) as shown in figure 4c.

**Supplementary figure 16.**
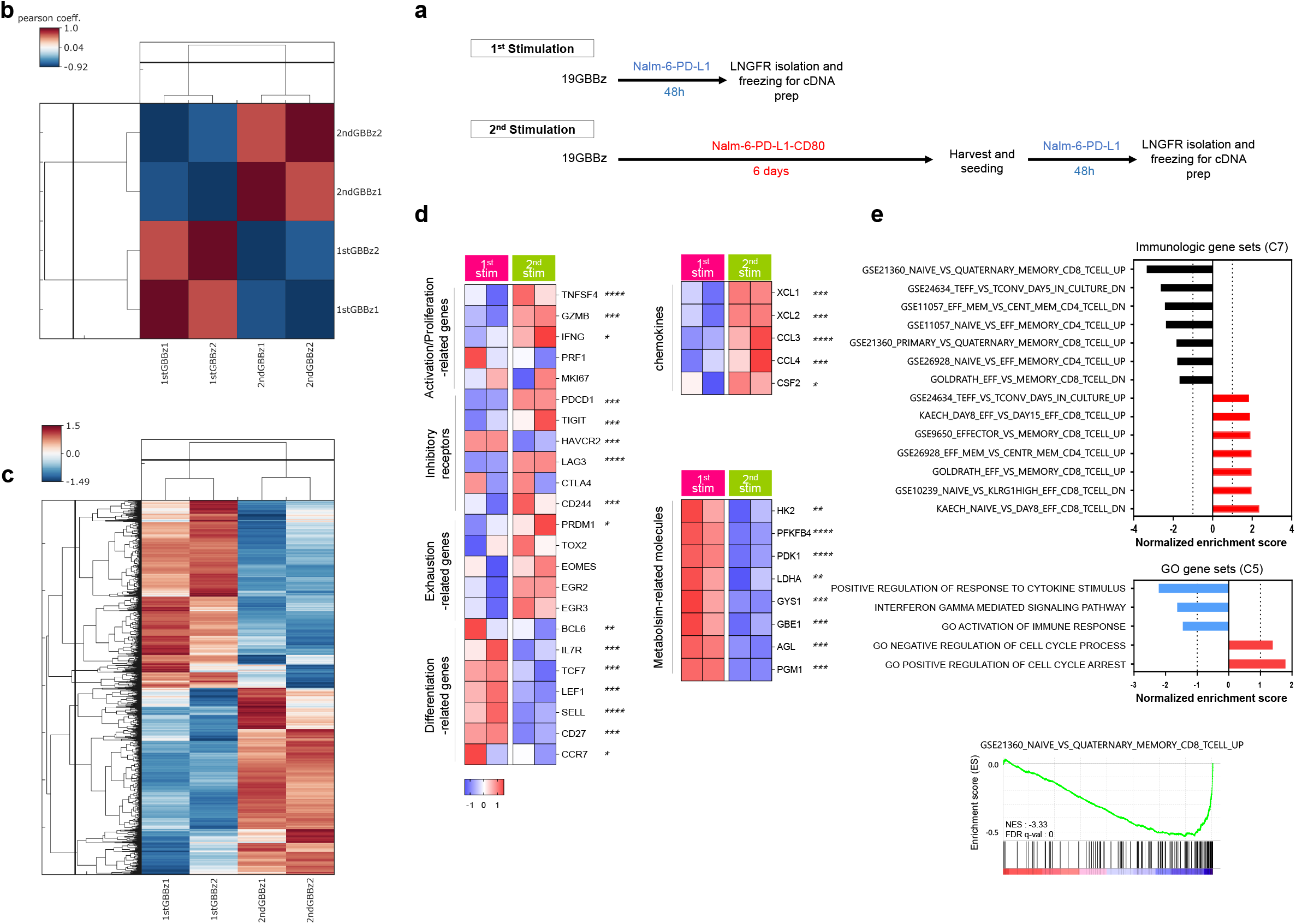
Changes in the transcriptomic profile of 1^st^ and 2^nd^ stimulated CAR T cells. **(a)** Schematic representation of the timeline for 1^st^ and 2^nd^ stimulation and sample preparation for RNA-seq. **(b)** Pearson’s correlation analysis of the transcriptomic profiles and **(c)** hierarchical clustering of differentially-expressed genes from 1^st^- and 2^nd^-stimulated 19GBBz cells derived from two donors (FDR *q ≤* 0.05). **(d)** Heat map of selected genes associated with T cell function in 1^st^- and 2^nd^-stimulated 19GBBz cells. Asterisks represent the statistical significance as measured by *q*-value. **(e)** Normalized Enrichment Scores (NESs) of significantly enriched gene sets associated with phenotypic and functional T cell signatures in 1^st^- and 2^nd^-stimulated 19GBBz cells as determined by GSEA analysis. * = *q* < 0.05, ** = *q* < 0.01, *** = *q* < 0.001, **** = *q* < 0.0001.

**Supplementary figure 17.**
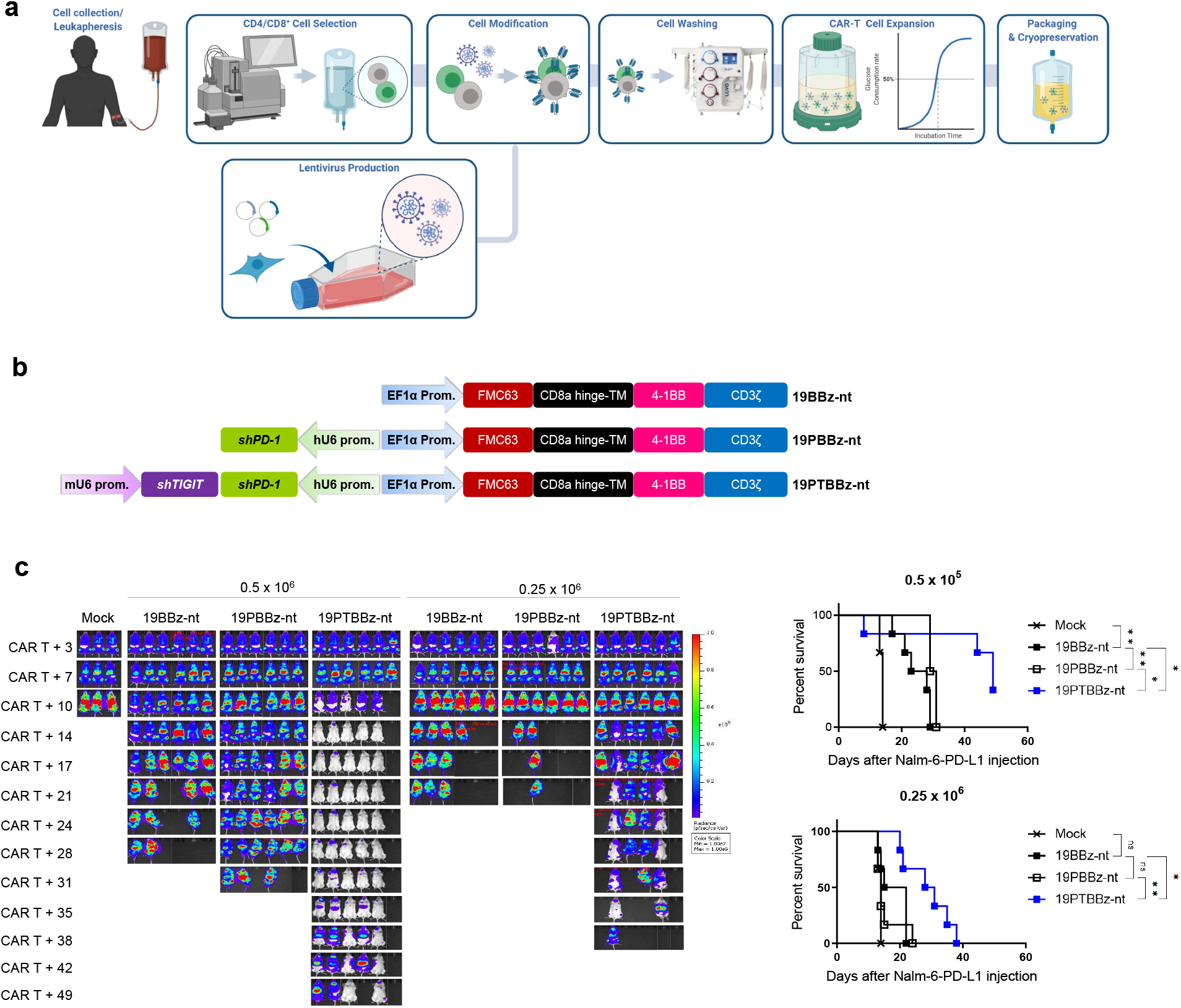
Generation and functional evaluation of healthy donor-derived, clinical-grade CD19-targeting CAR T cells. **(a)** Schematic representation of CAR T cell manufacturing in a semi-automated closed system. **(b)** Schematic representation of the vectors used to manufacture healthy donor-derived, clinical-grade CAR T cells. **(c)** NOG mice were injected intravenously with 1×10^6^ Nalm-6-GL-PD-L1 cells. 5 days later, CAR^+^ T cells (% CAR^+^: 19BBz-nt 40.19 %; 19PBBz-nt 38.46 %; 19PTBBz-nt 42.10 %) were intravenously injected at the indicated doses. Tumor burden was monitored based on the bioluminescence intensity from the IVIS imaging system. Kaplan-Meier survival analysis with Log-rank (Mantel-Cox) test comparing each CAR T-treated group. Data are from n = 3 mock mice and n = 6 mice for all CAR T cell-treated groups.

**Supplementary figure 18.**
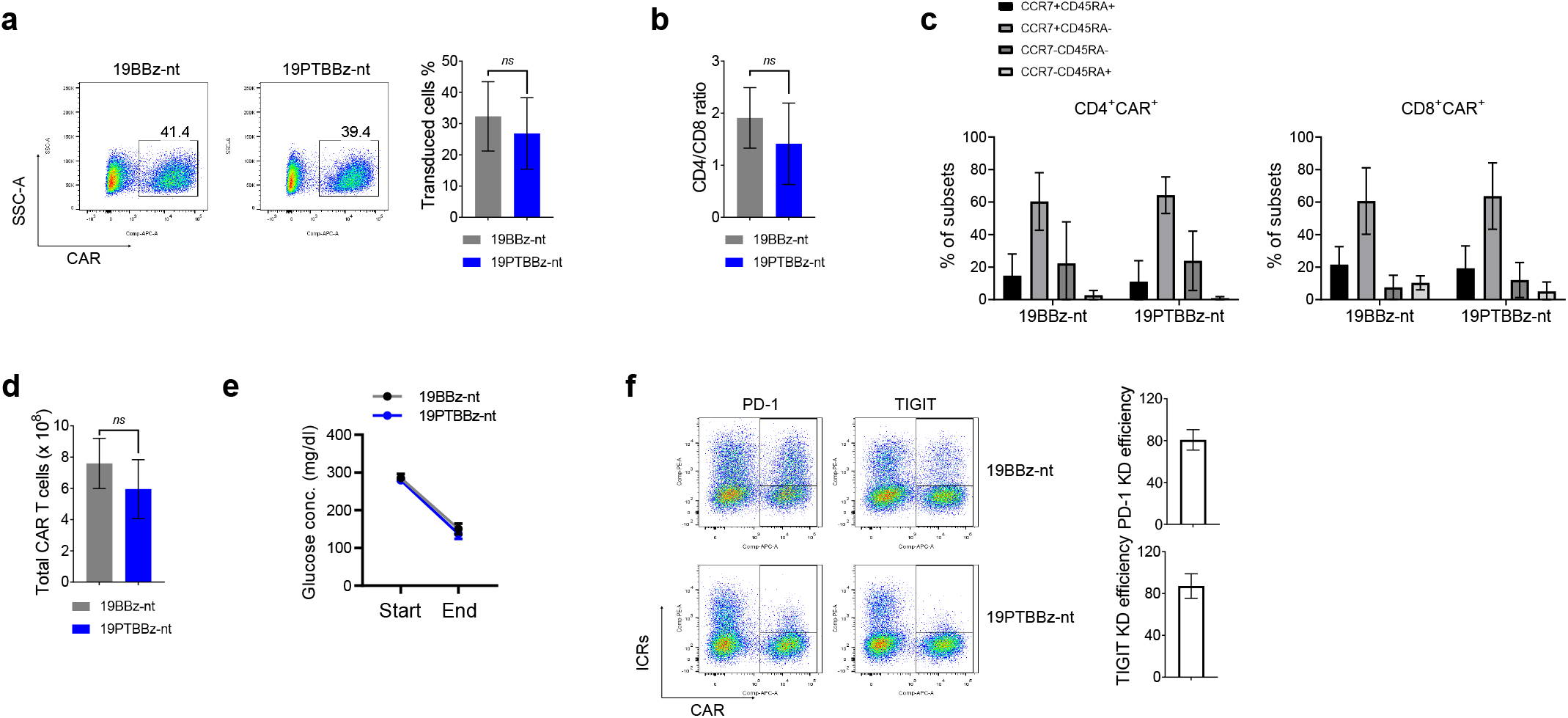
Generation and characterization of patient-derived, clinical-grade CD19-targeting CAR T cells with dual PD-1/TIGIT downregulation. **(a)** Transduction efficiency of CAR^+^ T cells on day 6 after transduction. **(b)** CD4/CD8 ratio of 19BBz-nt or 19PTBBz-nt cells on day 6 after transduction. **(c)** Expression of CD45RA and CCR7 to distinguish the differentiation state of 19BBz-nt or 19PTBBz-nt cells. **(d)** The absolute number of CAR T cells on day 6 after transduction. **(e)** Change in glucose concentration during the rapid expansion with the G-rex gas permeable culture device. **(f)** Knockdown efficiency of PD-1 and TIGIT. All data are the mean ± SD from three independent experiments. Statistical analysis was done by unpaired two-tailed t-test. T_N_ = CD45RA^+^CCR7^+^, T_CM_ = CD45RA^-^CCR7^+^, T_EM_ = CD45RA^-^CCR7^-^, T_EFF_ = CD45RA^+^CCR7^-^. nt = not tagged

## Materials and methods

### Cell lines and culture conditions

Nalm-6-GL cells were kindly provided by R. Kochenderfer (National Cancer Institute). K562 cells were purchased from the American Type Culture Collection (ATCC). Raji-PD-L1 cells were purchased from InvivoGen (USA). K562 cells were engineered to express human CD19 by lentiviral transduction. Nalm-6-GL, or K562-CD19 cells were transduced to express human PD-L1 (Sino Biological; HG10084-UT cDNA subcloned into a lentiviral vector) to generate Nalm-6-GL-PD-L1 and K562-CD19-PD-L1 cells. Raji-PD-L1 cells were transduced to express firefly luciferase-green fluorescent protein (ffluc/GFP) to generate Raji-GL-PD-L1 cells. Nalm-6-GL-PD-L1 cells were transduced with human CD80 (human CD80 ORF referring to NM_005191.4 was cloned to lentiviral vector) to generate Nalm-6-GL-PD-L1-CD80. Lenti-X 293T packaging cells were obtained from Takara Bio (Japan). Raji, K562, and Nalm-6 cell lines were cultured in RPMI-1640 medium (Gibco) supplemented with 10% heat-inactivated fetal bovine serum (FBS, Gibco), 2 mM L-glutamine (Gibco), and 1% penicillin/streptomycin (Gibco) in a humidified incubator with a 5% CO_2_ atmosphere at 37°C. Lenti-X 293T cells were cultured Dulbecco’s modified Eagle medium (Gibco) supplemented with 10% heat-inactivated FBS, 2 mM L-glutamine, 0.1 mM non-essential amino acids (Gibco), 1 mM sodium pyruvate (Gibco), and 1% penicillin/streptomycin. Cell line authentication was performed by Korea Cell Line Bank based on criteria established by the International Cell Line Authentication Committee. None of the cell lines used in this research are included in the commonly misidentified cell lines registry. Mycoplasma contamination tests were conducted by the Laboratory Animal Resources Center of the Korea Research Institute of Bioscience and Biotechnology before use in *in vivo* experiments and resulted negative.

### Plasmid construction

To construct the lentiviral transfer vector encoding the CD19-specific CAR, the anti-CD19 scFv (FMC63) was fused by overlapping PCR to the CD8α spacer and transmembrane domains, the 4-1BB (CD137) or CD28 costimulatory domains, and the CD3ζ signaling domain. To isolate transduced T cells, truncated LNGFR (ΔLNGFR) was amplified from pMACS-ΔLNGFR (Miltenyi Biotec) and cloned into the construct with a P2A sequence immediately downstream of the EF-1α promoter to generate pLV-EF-1α-ΔLNGFR-P2A-CD19 CAR vector. shRNA expression cassettes were added upstream to the pLV-EF-1α-ΔLNGFR-P2A-CD19 CAR vector in the antisense direction under the Pol III promoter (mU6, hU6, or hH1). For dual immune checkpoint inhibition, two shRNA-expressing modules, respectively controlled by the mU6 and hU6 promoters, were cloned-in facing each other upstream of the of the EF-1α promoter. Immune checkpoint targeting siRNA or shRNA sequences are listed in **Table 1**. GFP-targeting shRNA was used as the negative control.

**Table 1:**
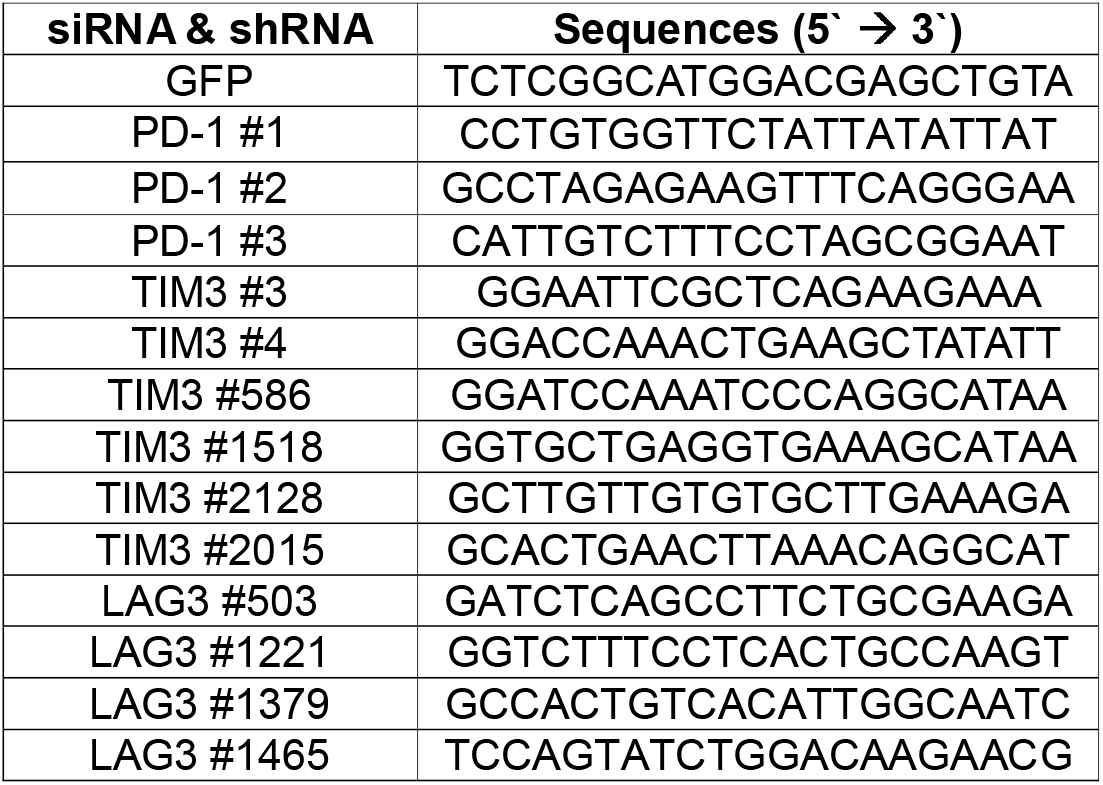
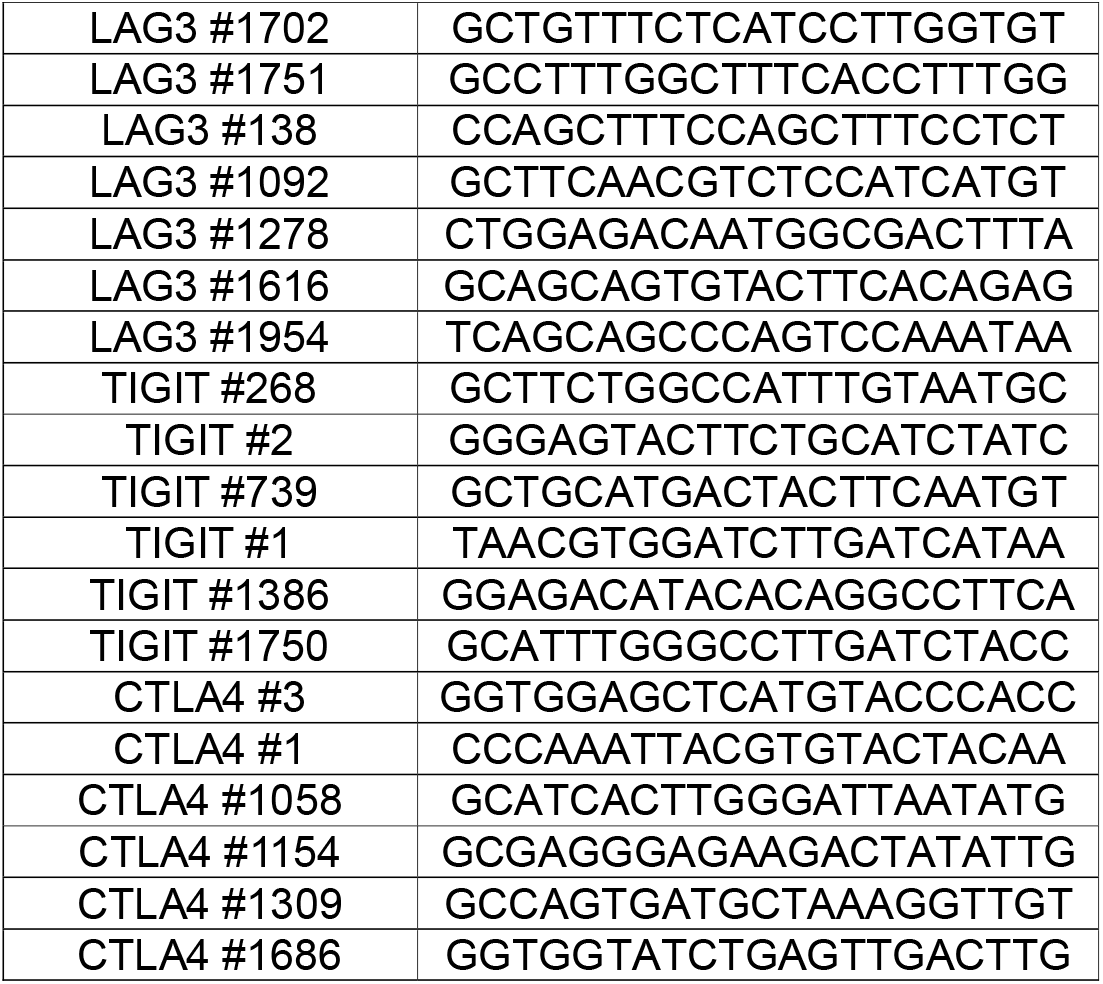
shRNA and siRNA sequences

### Generation of human CAR T cells.

Human PBMCs were obtained from healthy adult donors at the Seoul National University Hospital (SNUH) using protocols approved by the Institutional Review Boards (IRB number: H-1607-155-778). Lenti-X 293T cells were seeded at 6.5× 10^6^ cells per dish in Poly-D-Lysine-coated 100 mm dishes three days prior to transduction. After 24 hours, cells were co-transfected with 7.5 μg lentiviral transfer plasmid, 4.5 μg pMDG.1 encoding VSV-G envelope, 6 μg pRSV-Rev encoding Rev, and 6 μg pMDLg/pRRE encoding Gag/Pol using Lipofectamine 2000 according to the manufacturer’s instructions. Lentiviral supernatants were collected 40 hours following transfection, removed of cell debris by centrifugation at 1600 rpm for 5 min, and immediately used to transduce T cells. Peripheral blood mononuclear cells (PBMCs) were collected from whole blood samples of healthy donors using SepMate™ tubes (STEMCELL Technologies, Canada) in accordance with the manufacturer’s instructions. The PBMCs were stimulated with 4 μg/mL plate-bound anti-CD3 antibody (clone OKT3; Bio X cell), 2 μg/mL soluble anti-CD28 antibody (clone CD28.2; Bio X cell), and 300 IU/mL human recombinant IL-2 (BMI KOREA, Republic of Korea) in complete T cell medium containing 90% RPMI-1640 supplemented with 10% heat-inactivated FBS, 0.WmM non-essential amino acids, 2 mM GlutaMAX (Gibco, USA), 1mM sodium pyruvate, and 0.05 mM 2-mercaptoethanol (Gibco, USA). Two days after stimulation, activated T cells were transduced with lentiviral supernatants and 10 μg/ml protamine sulfate followed by centrifugation at 1000 × *g* for 90 min at 32°C and further incubated at 37°C. 24 hours later, the lentiviral supernatants were removed and the transduced T cells were expanded in the conditions described above. Fresh medium was added on day 3. The percentage of transduced T cells was evaluated by ΔLNGFR expression on day 4 after transduction. ΔLNGFR-positive transduced T cells were isolated using the human CD271 MicroBead kit (Cat# 130-099-023, Miltenyi Biotec) according to the manufacturer’s instructions. CAR^+^ ΔLNGFR^+^ T cells were maintained in complete T cell medium with 300 IU/mL rhIL-2.

### Generation of patient-derived, clinical-grade CAR T cells

The collection of human T cells from the leukapheresis product of a DLBCL patient (F, 54 years old) with recurrent brain metastasis who was administered 5 cycle of rituximab, MTX, Vincristine, and procavaine, was approved by the Institutional Review Board of the Samsung Medical Center (IRB #2018-11-066). A pool of CD4^+^ and CD8^+^ T cells was isolated using CliniMACS CD4 and CD8 GMP MicroBeads (Miltenyi Biotec, Germany). GMP-grade lentiviruses, generated with 19BBz-nt, 19PBBz-nt, or 19PTBBz-nt constructs, were manufactured by Takara Bio and used to generate CAR T cells per manufacturing protocols developed by Curocell Inc.

### Flow cytometry

All flow cytometry were performed with the LSRFortessa™ X-20 cytometer (BD, USA) and analyzed with the FlowJo software (Tree Star, USA). For cell surface staining, 2×10^5^ cells were stained with antibodies resuspended in 100 μL FACS buffer (1% BSA in DPBS) for 20 min at 4°C in the dark. Cells were washed with 2mL FACS buffer, resuspended, and analyzed. For intracellular staining of CTLA-4, T cells were fixed/permeabilized using 200 μL BD Cytofix/Cytoperm solution (BD, USA) for 20 min at 4°C. Cells were washed with 2mL intracellular staining buffer (1% BSA, 0.1% sodium azide, and 0.1% saponin in DPBS) and then stained with antibodies. Intracellular staining of IL-2 followed the same protocol as that for CTLA-4 but included the addition of 1 μg/mL GolgiPlug (BD Biosciences) to the culture media 12 hours prior to sample preparation. Viable cells were determined with Fixable Viability Dye eFluor™ 780 (ThermoFisher, USA). CD19-CAR moiety was labeled by AF647-conjugated anti-mouse F(ab’)_2_ antibody or biotin-conjugated rhCD19-Fc (ACRO Biosystems, USA) with AF647-conjugated streptavidin (Biolegend, USA). ΔLNGFR was labeled with APC-conjugated LNGFR antibody (clone ME20.4-1.H4, Miltenyi Biotec, Germany). All flow cytometry analyses were assessed using the following antibody clones. From BioLegend: CD4-BV605 (clone OKT4), CD8-APC or PE (clone SK1), CCR7-PE (Clone G034H7), CD45RO-PerCp-Cy5.5 (clone UCHL1), CTLA-4-PE (clone BNI3), LAG3-PE (clone 7H2C65), CD226-PE (clone 11A8), CD155-PE (clone SKII.4), CD112-PE (clone TX31), CD19-APC (clone HIB19), PD-L1-APC (clone 29E.2A3), CD80-PE (clone 2D10), HLA-DR-PE (clone L243), Galectin-9-PerCp-Cy5.5 (clone 9M1-3), and IL-2-PE (clone MQ1-17H12). From BD: CCR7-BV421 (clone 150503) and CD45RA-FITC (clone HI100). From Thermo: PD-1-PE (clone J105), TIGIT-PE (clone MBSA43). From R&D Systems: TIM3-PE (clone 344823)

### siRNA transfection

Stimulated T cells were transfected with 150 pmol siRNA at 1600 V for 10 ms with three pulses using the NEON transfection system (ThermoFisher). 24 hours later, knock-down efficiency was determined by flow cytometry. siRNA sequences, which can be converted shRNA, were listed in **Table 1**. In the case of PD-1 and TIM-3, however, the shRNA screening process proceeded immediately, skipping the selection of optimal siRNA candidates.

### PD-1/PD-L1-dependent proliferation assay

To determine the impact of the PD-L1/PD-1 axis on their proliferation, 1×10^6^ CAR T cells were co-cultured with γ-irradiated K562-CD19 or K562-CD19-PD-L1 at 1:3 effector:target (E:T) ratio for 7 days and counted with the countess II Automated Cell Counter (ThermoFisher, USA).

### Proliferation by repeated antigen stimulation

For 1^st^-stimulation, 1×10^6^ CAR T cells were co-cultured with γ-irradiated Nalm-6-GL-PD-L1-CD80 or K562-CD19-PD-L1 at 1:3 effector: target (E:T) ratio on day 0 for 6 days. For second stimulation, CAR T cells were harvested and numerated with the countess II Automated Cell Counter before re-seeding (1×10^6^ cells) with the same target cells at a 1:3 E:T ratio for another 6 days before further cell counting. To evaluate the contribution of CD226 signaling to the proliferative activity of 19PTBBz cells, day 1^st^-stimulated CAR T cells were co-cultured with γ-irradiated target cells at a 1:3 E:T ratio in the presence of 10 μg/ml plate-bound human CD226 blocking monoclonal antibody (clone DX11, Abcam, UK). Ultra-LEAF™ purified mouse IgG1(Clone MOPC-21, Biolegend, USA) was used as for isotype control.

### IncuCyte-based cytotoxicity

To evaluate the cytotoxic activity of CAR T cells in the presence or absence of PD-L1, 1×10^5^ Nalm-6-GL or Nalm-6-GL-PD-L1 cells were co-cultured with CAR T cells at a 1:1, 1:0.3, or 0.1:1 E:T ratio in 250 μL complete T cell medium in 96-well flat-bottom plates for up to 72 h. Triplicate wells were used for each CAR T group. GFP fluorescent intensity was detected every 2 hours using the IncuCyte S3 Live-Cell analysis system (Sartorius, Germany). Total integrated GFP intensity per well was used as a quantitative measure of viable target cells. Values of total integrated GFP intensity were normalized to the GFP intensity of the starting point.

### *in vivo* xenograft models

All animal testing procedures described here were approved by the Korea Advanced Institute of Science and Technology (KAIST) and Osong Medical Innovation Foundation Laboratory Animal Center Institutional Animal Care and Use Committee and were performed according to national and institutional guidelines. Immunocompromised NOD.Cg-Prkdc^scid^IL2γg^tm1Wjl^/JicKoat (NSG) mice were obtained from The Jackson Laboratory and bred in the KAIST Laboratory Animal Resource Center under a protocol approved by the KAIST Institutional Animal Care and Use Committee. 6-8 weeks-old female (NOD.Cg-Prkdc^scid^IL2γg^tm1Sug^/JicKoat) (NOG) mice were obtained from Koatech (Republic of Korea). 6-8 weeks-old NSG mice were administered with 1×10^6^ Nalm-6-GL or Nalm-6-GL-PD-L1 cells by tail vein injection. 5 days post-injection, the indicated dose of CAR T cells was injected intravenously. Tumor burden was measured after intraperitoneal injection of luciferin (PerkinElmer) per the manufacturer’s instructions and subsequent bioluminescence imaging using the IVIS Lumina X5 imaging system. Bioluminescence values were analyzed using the Living Image software (PerkinElmer). Tumor growth was measured weekly. To evaluate the *in vivo* efficacy of CAR T cell against a subcutaneous tumor model, 5×10^6^ Raji-GL-PD-L1 lymphoma cells were subcutaneously injected into the right flank of NOG mice. When the mean tumor volume reached approximately 100 mm^3^ (measured with a caliper), CAR T cells were intravenously injected into the mice at the indicated doses. Tumor growth was determined weekly by bioluminescence imaging. Absolute CAR T cell counts in the peripheral blood of mice were determined using CountBright Absolute Counting Beads (ThermoFisher, USA) according to the manufacturer’s instructions.

### CD266 knockout by Cas9 RNP

CD226 gRNA candidates, listed in **Table 2**, were predicted using the CHOPCHOP and Cas-Designer tools. The gRNA sequence, which contains the T7 promoter sequence, ~20 nucleotides of CD226-specific sequence and a gRNA backbone sequence, was overlapped with the sgRNA scaffold oligonucleotide by overlapping PCR. For *in vitro* transcription (IVT), 1.4 μg PCR products were mixed with 50U T7 RNA polymerase (NEB), 1 U RNase inhibitor (NEB), 10 mM fresh DTT, 14 mM MgCl_2_, and 4 mM of each ribonucleotide triphosphate (Jena Bioscience) and incubated at 37 °C overnight. For the eradication of DNA, 2 U DNase I (NEB) was added to IVT mixture and further incubated at 37 °C for 30 min. sgRNA was purified with the RNeasy MinElute Cleanup kit (Qiagen, 74204). sgRNA targeting the CAG (CMV-IE, chicken actin, rabbit beta globin) promoter was used as negative control. Cas9 RNPs were prepared immediately before electroporation by incubating 7 μg Cas9 (enzynomics, M058UL), and 14 μg synthetic sgRNA in 20 μM Hepes (pH 7.5), 150 mM KCl, 1 mM MgCl2, 10% (vol/vol) glycerol, and 1 mM TCEP at 37 °C for 10 min. Stimulated T cells were electroporated with the Neon™ transfection device and Transfection System 10 μl Kit (ThermoFisher, USA). 1.3×10^6^ cells were washed with PBS before resuspension in 8 μl T buffer. RNP and T cells were mixed and transferred to 10 μl Neon^TM^ Tip, and electroporated with a Neon electroporation device (1,600V, 10ms, 3 pulses). The electroporated T cells were suspended with fresh complete T cell medium, and further incubated with a 5% CO_2_ atmosphere at 37°C overnight. 24 hours post-electroporation, CD266 expression was evaluated by flow cytometry. Editing efficiency was estimated by T7 endonuclease I (NEB) assay according to the manufacturer’s instructions using the following primers: forward 5’-AAGTATCTTCCAGTTGGGTGTCCCA-3’, reverse 5’-GATTGATGGATTCACAAGATAAAGAAACCCTC-3’

**Table 2:**
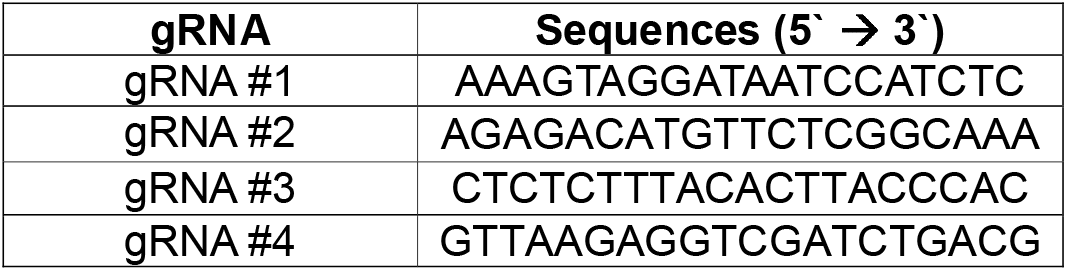
CD226-targeting gRNA sequences

### Bulk RNA-seq

1^st^-stimulated CAR T cells (2×10^7^) were prepared by culturing over γ-irradiated Nalm-6-PD-L1 cells for 48 hours prior to LNGFR isolation and freezing for cDNA preparation. 2^nd^-stimulated CAR T cells were prepared by culturing over γ-irradiated Nalm-6-PD-L1-CD80 cells for 6 days followed by harvesting and seeding 2×10^7^ cells over γ-irradiated Nalm-6-PD-L1 cells for 48 hours, LNGFR isolation, and freezing for cDNA preparation. Total RNA was isolated using the NucleoSpin RNA XS kit (MACHEREY-NAGEL, Germany) according to the manufacturer’s instructions. RNA quality was determined with the Agilent 4200 TapeStation (RIN value > 9). RNA sequencing libraries were constructed using the Illumina TruSeq Stranded mRNA LP kit. Briefly, mRNA was purified by oligo-dT beads and fragmented through enzymatic reaction. After fragmentation, cDNA was generated through reverse transcription. The cDNA libraries were constructed and followed by 100 bp paired-end mode on DNBSEQ-400 platform. External RNA controls consortium (ERCC) RNA spike-in mixes (Thermo Fisher, 4456740) were included for quality assurance. The quality of all paired-end reads was analyzed using FastQC software, and the per base sequence quality of all sample was above Q30. RNA-seq analysis was carried out via the Galaxy platform (https://usegalaxy.org/). FASTQ data was mapped to the reference genome hg19 using the HISAT2 v2.1.0 software. The number of reads per annotated gene was then computed from the mapped reads using the featureCounts v1.6.4 software. The R package limma with voom method v3.38.3 was used to normalize all data sets and analyze differential expression between groups. Pearson’s correlation matrices and hierarchical clustering plots of differentially expressed genes (FDR < 0.1) were generated with the Instant Clue software. The heatmap of selected genes associated with T cell function was generated using GraphPad prism 8. Each row represents the z-score of normalized gene expression values for the selected genes. Gene set enrichment analysis (GSEA) was performed using a preranked file generated by t-statistic on curated gene sets from the Broad Institute Molecular Signature Database (http://www.broadinstitute.org/gsea/msigdb/). Normalized data are accessible on a public database (GEO submission number GSE158676).

### Statistical analysis

Statistical analyses were determined by Student’s t-test (two-tailed, unpaired), one-way ANOVA, or two-way ANOVA. Percent survival was estimated using the Kaplan-Meier method, and statistical significance was calculated by the log-rank test. All statistical analyses were conducted using GraphPad Prism 8 (GraphPad Software, Inc). For all analyses, a p-value < 0.05 was considered statistically significant.

## ACKNOWLEDGEMENTS

This work was supported by Curocell Inc., by the Osong Medical Innovation foundation funded by the Ministry of Health & Welfare (2016M3A9D9945471, HO18C0005), by the National Research Foundation of Korea (NRF) (2017R1A6A3A11031455), by the Korea Health Technology R&D Project through the Korea Health Industry Development Institute (KHIDI), by the Ministry of Health & Welfare (HI20C0043), and by the KAIST Global Centre for Open Research with Enterprise (GCORE) grant funded from the Ministry of Science and ICT (N11190028).

## AUTHOR CONTRIBUTIONS

CHK, YHL, and HCK designed the experiments and wrote the manuscript. YHL, HJL, YJL, SKN, JSM, XW, and OS performed the experiments and analyzed the data. CH analyzed the data and wrote the manuscript. WSK, SJK, YK, and IJ designed the experiments and analyzed the data.

